# Dimerization-dependent *cis*-autophosphorylation activates the UPR and ISR kinases

**DOI:** 10.64898/2026.07.28.741179

**Authors:** Magdalena Otto, Thomas Leonard

## Abstract

Maintaining cellular homeostasis requires the ability to detect and respond to various forms of stress. Protein kinases of the unfolded protein response (UPR) monitor protein folding in the endoplasmic reticulum, while the integrated stress response (ISR) kinases sense viral infection, nutrient deprivation, mitochondrial oxidative stress and proteotoxic stress. However, the precise molecular mechanism by which these kinases are activated is still unknown. In this study, we show that the ISR kinases PERK, HRI, and Gcn2, as well as the UPR kinase Ire1, undergo dimerization-dependent *cis*-autophosphorylation of their activation loops. The back-to-back dimerization of the kinase domain allosterically activates each kinase. We derive a simple mathematical model for dimerization-dependent *cis*-autophosphorylation, which we use to obtain kinetic parameters of PERK autophosphorylation. We show that dimerization promotes not only activation loop phosphorylation of PERK, but also phosphorylation of its substrate, eIF2α. In cells, kinase domain dimerization is necessary and sufficient for the activation of PERK. In summary, we propose a model in which a dimer is the minimal functional unit of all UPR and ISR kinases.

## Introduction

Cells are constantly exposed to various forms of intracellular stress. Diverse stress signals are transduced by the integrated stress response (ISR), which is mediated by four kinases to mount adaptive transcriptional and translational responses to restore cellular homeostasis. Protein kinase RNA (PKR)-like ER kinase (PERK) senses unfolded proteins in the lumen of the ER^1^, while General Control Nonderepressible 2 kinase (Gcn2) plays a role in sensing unloaded tRNA (amino acid starvation)^2,3^. Heme-regulated inhibitor kinase (HRI) is a heme sensor in erythroid progenitors and has recently been shown to sense mitochondrial stress^4^ and trigger mitophagy^5^, while protein kinase R (PKR) plays a role in anti-viral defense^6^. Dysregulation of ISR signaling has been linked to inflammation, cancer and neurodegenerative disease^7^ and inhibition of the ISR has been shown to have a neuroprotective effect by preventing stress granule formation^8,9^.

The ISR kinases all converge on the same substrate, phosphorylating the eukaryotic translation initiation factor 2α (eIF2α) on S52^1,10–12^. eIF2α is part of the tripartite eIF2 complex, a translation initiation factor which recruits the Met-tRNA_i_^Met^ to the small ribosomal subunit in a GTP-dependent manner. Phosphorylation of eIF2α turns eIF2 into an inhibitor of its guanine nucleotide exchange factor (GEF), eIF2B, reducing translation initiation and hence decreasing protein folding load^13–15^.

Surveillance of protein folding load is especially important in the endoplasmic reticulum (ER), which handles around a third of the cell’s proteome^16^. The accumulation of unfolded proteins in the ER triggers the Unfolded Protein Response (UPR). The UPR is orchestrated by three transmembrane proteins, activating transcription factor-6 (ATF6) and two kinases: inositol-requiring protein-1 (Ire1) and PERK^17^. Mutations in *EIF2AK3*, the gene encoding PERK, cause Wolcott-Rallison Syndrome, a disease characterized by permanent neonatal or early infancy insulin-dependent diabetes^18^, while PERK knockout mice exhibit early onset *Diabetes mellitus* due to impairment of insulin secretion in the pancreas^19^. Ire1 is essential for metazoan embryonic development^20^ and dysregulation of Ire1 is associated with pro-apoptotic phenotypes^21^.

Among the UPR sensors, Ire1 and PERK share a conserved ER luminal domain, which senses unfolded proteins in the ER. Both solution data and crystal structures of the core luminal domains of human and yeast Ire1 as well as human PERK show a constitutively dimeric assembly^22–24^, and the ISR kinases Gcn2, HRI and PKR have all been reported to dimerize via their respective sensory domains^12,25–27^. The sensory domains of each protein kinase transduce the respective stress signal into activation of the kinase domain via autophosphorylation of its activation loop^28–32^. Autophosphorylation of the ISR kinases triggers phosphorylation of eIF2α^1,10–12^, while autophosphorylation of Ire1 has been proposed to activate its cytoplasmic RNase domain^33^.

Crystal structures of the kinase domains of PERK, PKR, Gcn2 and Ire1 have all revealed their dimerization in a back-to-back configuration^30,34–37^, a conformation which is incompatible with the canonical mechanism of activation loop phosphorylation in *trans* (intermolecular). Although PKR has been described to autophosphorylate in *cis* (intramolecular)^38^, Gcn2 and HRI have been described to *trans*-autophosphorylate^32,39^. The prevailing model for Ire1 autophosphorylation depends on the formation of higher-order oligomers, which have been proposed to autophosphorylate adjacent dimers in *trans*^37^. The oligomerization of Ire1 and PERK is hypothesized to be mediated by their luminal domains^22,23,37,40^, which are functionally interchangeable^41^, thereby driving clustering in the ER membrane^42–47^. The filamentous arrangement of Ire1 protomers in a crystal structure of the Ire1 kinase and RNase domains has therefore been suggested to represent the active conformation^37^. The structure of a face-to-face arrangement of Ire1 has since been proposed to represent the *trans*-autophosphorylation reaction via activation loop exchange^28^. Due to the structural homology of the PERK and Ire1 luminal domains, oligomerization-mediated *trans*-autophosphorylation has also been proposed for PERK^1,48^. Despite these observations, however, several factors argue against this model, including the low nanomolar concentrations of these proteins in the cell^49^, significant barriers to diffusion in the membrane^50,51^, and face-to-face dimerization, which, by definition, is sterically incompatible with subsequent substrate phosphorylation.

In this study, we show that the minimal functional unit of the UPR and ISR kinases is a dimer. We demonstrate that PERK undergoes dimerization-dependent *cis*-autophosphorylation and that the dimeric conformation of PERK drives eIF2α phosphorylation. We present a kinetic model for dimerization-dependent activation loop *cis*-autophosphorylation and show that dimerization is necessary and sufficient to activate PERK in cells. Finally, we show that dimerization-dependent *cis*-autophosphorylation activates all of the UPR and ISR kinases.

## Results

### Dimerization of PERK is sufficient to drive activation loop *cis*-autophosphorylation

To establish whether PERK is capable of activation loop autophosphorylation in *cis*, we purified a recombinant PERK kinase domain construct (PERK^KD^). To prevent off-target phosphorylation of sites other than the canonical activation loop T980, we purified a construct of the PERK kinase domain that was previously crystallized (Figure 1A, PDB: 3QD2^30^) and iteratively removed or mutated residues which were non-specifically phosphorylated *in vitro* (Supplementary Figure 1, Supplementary Table 1). Tandem mass spectrometry revealed that these residues were all located in an insertion loop of the N-lobe of the kinase (Figure 1A, Supplementary Figure 1) which has previously been shown not to affect activation loop autophosphorylation^52^.

**Figure 1.**
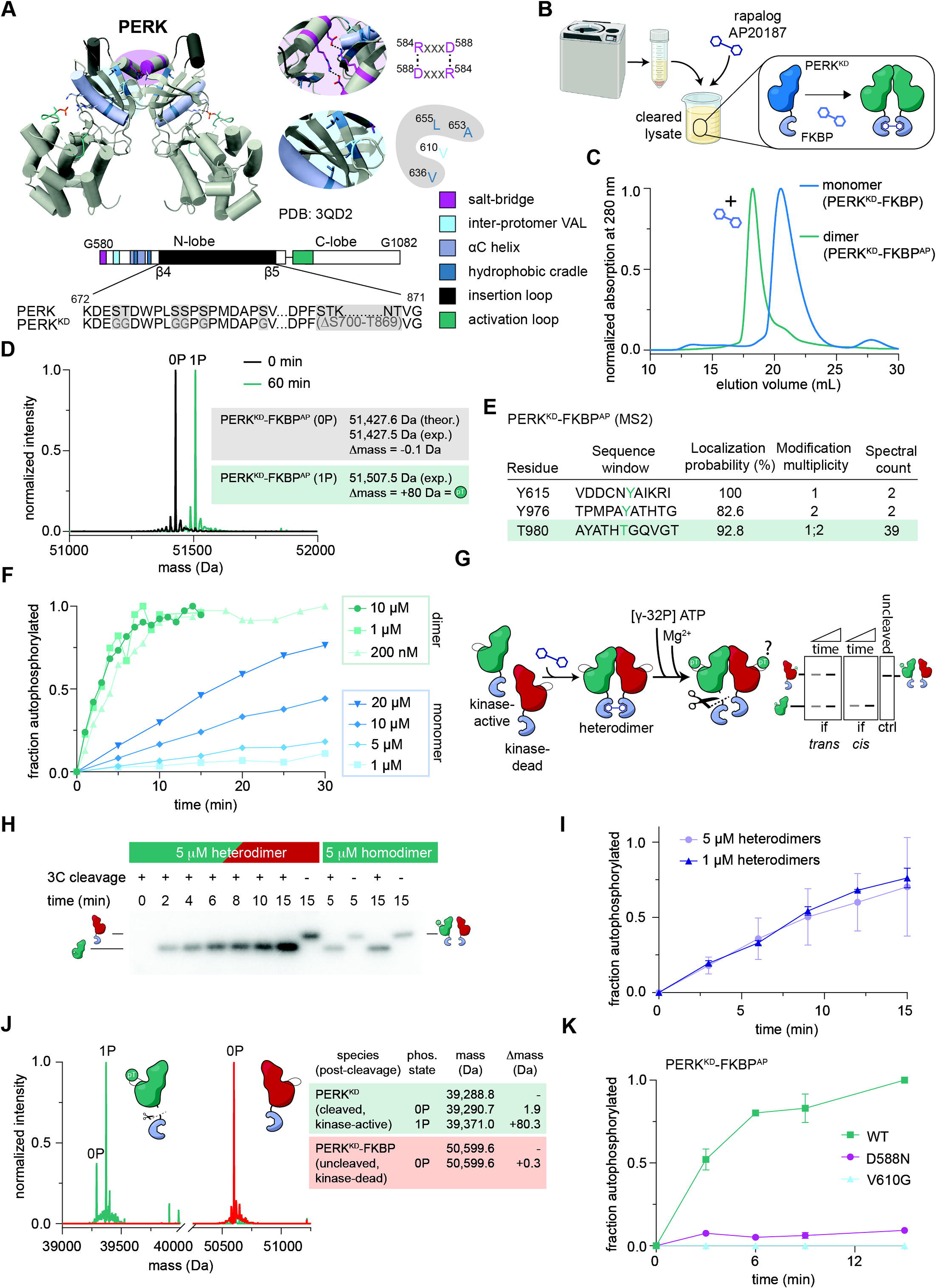
Dimerization of PERK is sufficient to drive activation loop *cis*-autophosphorylation. A. Crystal structure of the active, dimeric PERK kinase domain phosphorylated on T980 (PDB: 3QD2). Relevant features for catalysis and dimerization in the kinase are highlighted in the structure and domain architecture (magenta: inter-dimer salt bridge, light blue: inter-protomer valine (VAL), violet: α-C helix, dark blue: hydrophobic cradle, black: insertion loop in the kinase between the β4 and β5 sheets, green: activation loop). With reference to the wildtype protein (PERK), all mutations and deletions introduced in the insertion loop are depicted in PERK^KD^. B. Chemically-induced dimerization of PERK^KD^-FKBP (violet). Upon addition of the dimerizer AP20187 to the cleared lysate, monomeric PERK^KD^-FKBP (blue) dimerizes (green). C. Size-exclusion chromatogram of PERK^KD^-FKBP -AP20187 (blue) and +AP20187 (green). D. Intact mass spectrometry of PERK^KD^-FKBP^AP^ after 0 minutes and 60 minutes of autophosphorylation. E. Tandem MS analysis of PERK^KD^-FKBP^AP^ to identify phosphosites after 60 minutes of autophosphorylation (Figure 1D). F. Time course of radiometric autophosphorylation assay at different concentrations of PERK^KD-^FKBP (blue) and PERK^KD^-FKBP^AP^ (green). G. Experimental strategy to distinguish between *cis* and *trans*-autophosphorylation. PERK heterodimers comprising kinase-active and kinase-inactive protomers were autophosphorylated *in vitro* followed by proteolytic cleavage to distinguish the two protomers by SDS-PAGE. Predicted outcomes are depicted for both *cis* and *trans* reactions. H. Representative phosphorimage of radiometric autophosphorylation time course of cleaved PERK heterodimers. I. Quantification of time course of autophosphorylation PERK^KD^ heterodimers at 1 μM and 5 μM. J. Intact mass spectrometry of PERK^KD^ heterodimers after 1 h autophosphorylation at 1 μM. K. Time course of radiometric autophosphorylation assay of 1 μM wildtype PERK^KD^-FKBP^AP^ and interface mutant (D588N, V610G) dimers.

In order to compare the autophosphorylation kinetics of monomers and dimers, we next established an *in vitro* dimerization system (Figure 1B). Dimerization systems (constitutive and chemically inducible) have been extensively employed to investigate the UPR and ISR kinases in cells^38,53–55^, but not *in vitro*. Fusion of the FKBP12^F36V^ homo-dimerization domain^56^ to the C-terminus of PERK^KD^ (PERK^KD^-FKBP) and addition of the dimerizer AP20187 to the cleared cell lysate resulted in a shift in the size exclusion chromatogram of the purified protein to a higher molecular weight (Figure 1C). AP20187-mediated dimerization of PERK^KD^-FKBP (henceforth denoted by PERK^KD^-FKBP^AP^) was confirmed by mass photometry, with an apparent dissociation constant of 3.4 nM (Supplementary Figure 2A), consistent with the reported affinity of FKBP for the dimerizer of 1.8 nM^56^. We confirmed that PERK^KD^-FKBP^AP^ was completely dephosphorylated during purification and stoichiometrically phosphorylated exclusively on T980 of its activation loop after a 60 min autophosphorylation assay (Figure 1D-E).

To examine the impact of dimerization on the kinetics of T980 autophosphorylation, we subjected both monomers and dimers of PERK^KD^-FKBP to kinase assays. Monomeric PERK^KD^-FKBP exhibited a concentration-dependent increase in the rate of autophosphorylation, whereas autophosphorylation of dimeric PERK^KD^-FKBP^AP^ exhibited concentration-independent kinetics (Figure 1F). These observations indicate that autophosphorylation likely does not occur in *trans* between dimers, but as an intramolecular *cis* reaction stimulated, in *trans,* by dimerization.

To differentiate between *trans*-autophosphorylation and dimerization-dependent *cis*-autophosphorylation, we purified heterodimers comprising kinase-active and kinase-dead (D935N) protomers. Following autophosphorylation, the active protomer was cleaved with 3C protease to generate distinct fragments with different molecular weights that could be distinguished by gel electrophoresis (Figure 1G, Supplementary Figure 3A). While autophosphorylation of the kinase-active protomer increased over time, there was no signal for the kinase-dead protomer (Figure 1H), demonstrating that no *trans* reaction occurred in this assay. The uncleaved heterodimer was loaded alongside the time course as a control, as both protomers run with the same electrophoretic mobility of the non-cleavable, kinase-inactive protomer. The autophosphorylation rate of the heterodimers was concentration-independent (Figure 1I), as already shown for kinase-active homodimers (Figure 1F), consistent with a *cis* reaction. Mass spectrometry analysis of the reaction products confirmed that only the kinase-active protomer was mono-phosphorylated (Figure 1J), confirming that the reaction occurred exclusively in *cis*.

Having established that activation loop autophosphorylation occurs in *cis*, we next investigated how dimerization allosterically drives PERK autophosphorylation. The crystal structure of the PERK kinase domain^30^ reveals two conserved features: reciprocal salt bridges on the N-lobe between R584 and D588, and the insertion of V610 from one protomer into a hydrophobic pocket formed by V636, A653 and L655 of the other protomer (Figure 1A). We introduced mutations in both the salt bridge (D588N) as well as the hydrophobic residue (V610G) to disrupt inter-protomer contacts. We confirmed their expected mass by mass spectrometry (Supplementary Figure 3B-C) and verified protein folding by nano differential scanning fluorimetry and circular dichroism spectroscopy (Supplementary Figures 4A and 4B). Using in-solution crosslinking, we confirmed that the mutations impaired dimerization as predicted (Supplementary Figure 4C). In the context of dimeric PERK^KD^-FKBP^AP^, mutation of the R584-D588 salt bridge significantly decreased the rate of autophosphorylation, while removal of the hydrophobic side chain of V610 completely abrogated autophosphorylation (Figure 1K). These observations imply that specific inter-protomer contacts are required for allosteric activation that cannot be overridden by forced dimerization.

In summary, PERK dimerization is both necessary and sufficient for stoichiometric activation loop *cis*-autophosphorylation in vitro.

### Kinetic model of dimerization-dependent *cis*-autophosphorylation

To obtain insights into the dimerization-dependent *cis*-autophosphorylation reaction mechanism, we modeled its kinetics. Dimerization is governed by the association and dissociation rate constants *k_a_* and *k_d_*, respectively, while phosphorylation can be described by the rate constant *k_cat_* (Figure 2A). The reverse reaction (dephosphorylation) is considered to be zero. To model the kinetics of autophosphorylation with as few parameters as possible, we made two assumptions: (i) the affinity of homodimerization (*K_D_*) is not affected by phosphorylation and (ii) the catalytic rate constants of symmetric and asymmetric autophosphorylation are equal (Figure 2A).

**Figure 2.**
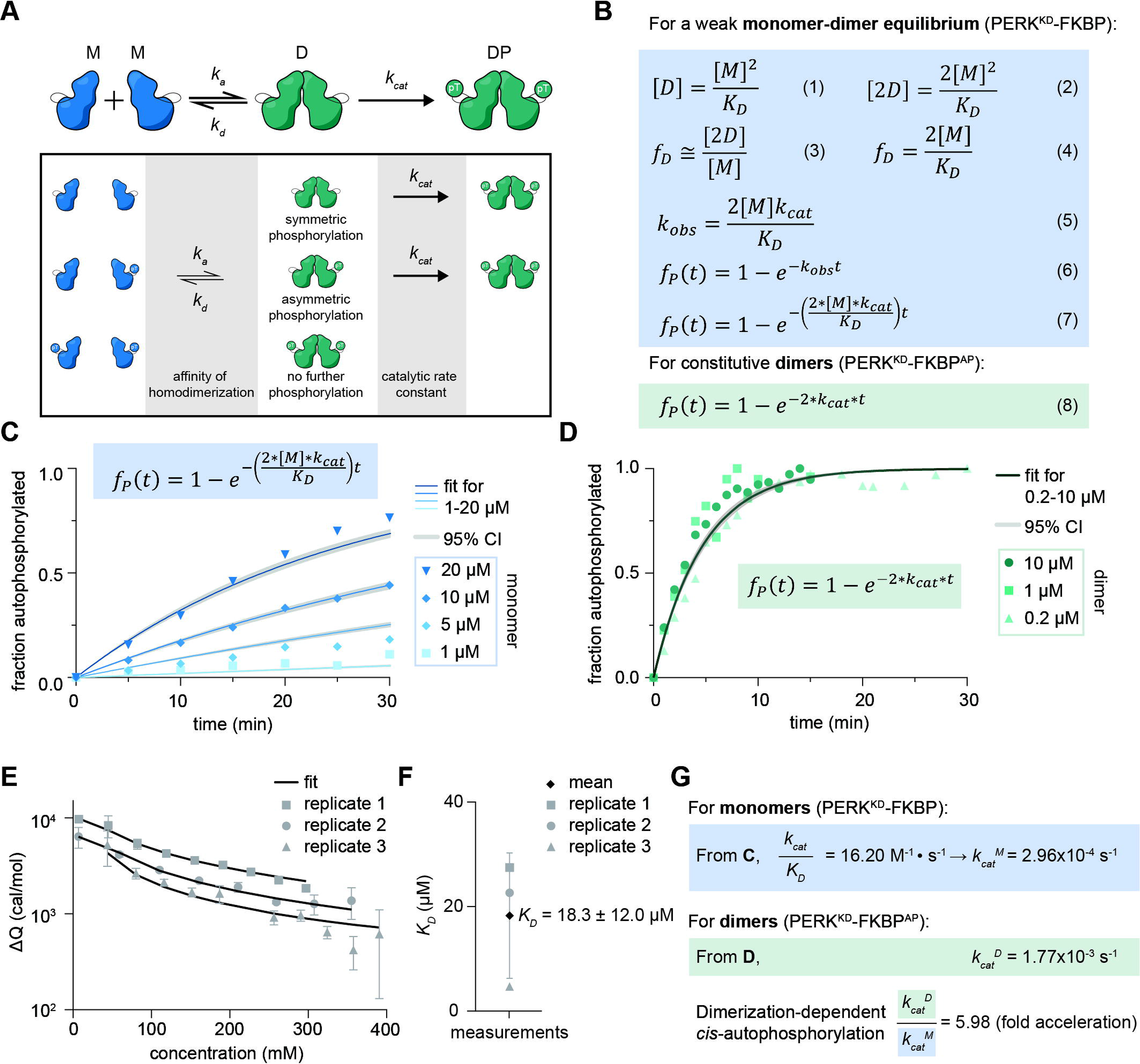
Kinetic model of *trans*-activated *cis*-autophosphorylation. A. Model of back-to-back *cis*-autophosphorylation, including parameters and assumptions of the model. M= monomeric kinase, D = dimeric kinase, DP = dimeric phosphorylated kinase B. Equations and derivations to mathematically model the rate of autophosphorylation. C. PERK^KD^-FKBP autophosphorylation curves with global fits to equation (7). Fits for 4 different concentrations are shown as solid lines and the corresponding 95% confidence intervals (CI) are displayed in grey. D. PERK^KD^-FKBP^AP^ autophosphorylation curves with global fits to equation (8). The fit is displayed as a solid line with the corresponding 95% confidence interval (CI) in grey. E. Isothermal titration calorimetry of PERK^KD^ (n = 3 independent injection series). Data were globally fit with a one-sided dissociation model. F. *K_D_* of PERK^KD^ homodimerization derived from the fit to each injection series in E. G. Derivation of *k_cat_* for monomers (PERK^KD^-FKBP) and dimers (PERK^KD^-FKBP^AP^).

The concentration of dimers [D] in an equilibrated solution of monomers [M] and dimers depends on the affinity of homodimerization, *K_D_,* and is given by equation (1) (Figure 2B, Supplementary Figure 5A and 5B). Since only dimers are phosphorylation-competent, the concentration of phosphorylation-competent molecules in an equilibrated solution is defined by equation (2). For a weak monomer-dimer equilibrium, the fraction of dimeric molecules, *f_D_*, is approximated by equation (3). Substituting (2) into (3) defines *f_D_* in terms of monomer concentration and affinity of homodimerization. The pseudo-first order rate of autophosphorylation observed experimentally (*k_obs_*) is defined by equation (5) (Supplementary Figure 5, catalytic rate constant) under the saturating ATP concentrations of our experimental setup. Equation (6) describes the accumulation of phosphorylated kinase as a function of time (Supplementary Figure 5, equations (6) and (7)). Substituting (5) into (6) gives equation (7), in which the autophosphorylation kinetics are described in terms of *k_cat_*, *K_D_* and [M]. For PERK^KD^-FKBP^AP^, in which dimerization is constitutive, we can simplify the reaction kinetics with equation (8), since the reaction is concentration-independent.

Using equations (7) and (8), we applied a global fit of the parameter *k_cat_*/*K_D_* for the monomer curves (Figure 2C) and of parameter *k_cat_* for the dimer curves (Figure 2D), respectively. A global fit to all monomer and dimer datasets is appropriate because *k_cat_*and *K_D_* are constants. To determine *k_cat_* of the reaction for the monomer, we determined the *K_D_* of homodimerization using isothermal titration calorimetry. We injected concentrated PERK^KD^ into buffer and measured the raw heat (ΔQ) of the injection (Figure 2E). By fitting a global dissociation model (Supplementary Figure 6) to three independent injection series, we obtained a value for *K_D_* of 18.3 ± 12.0 μM (Figure 2F), yielding a monomeric *k_cat_* (*k_cat_^M^*) of 2.96*10^-4^ s^-1^ (Figure 2G). Fitting equation (8) to the PERK^KD^-FKBP^AP^ autophosphorylation curves, we obtained a value for *k_cat_^D^* of 1.77*10^-3^ s^-1^ (Figure 2G), corresponding to a 6-fold increase in catalytic rate upon dimerization.

In summary, the kinetics of dimerization-dependent *cis*-autophosphorylation can be quantitatively modeled with a simple equation that links two intrinsic properties of the PERK kinase domain: its homodimerization affinity (K_D_) and its catalytic rate constant (k_Cat_).

### Dimerization is sufficient for PERK activation in cells

Dimerization is necessary and sufficient to activate PERK *in vitro*. To test this hypothesis directly in cells we again turned to the FKBP dimerization system. We expressed a C-terminal FKBP fusion of full-length PERK in mammalian HEK293 cells. It has already been established that overexpression of PERK leads to its autophosphorylation and activation of the UPR^1,57^. Thus, to control for hyperactivation of the pathway induced by overexpression, we compared two promoters of different strength: the HSV-TK promoter (near endogenous expression^58^) and the CMV promoter (overexpression) (Figure 3A).

**Figure 3.**
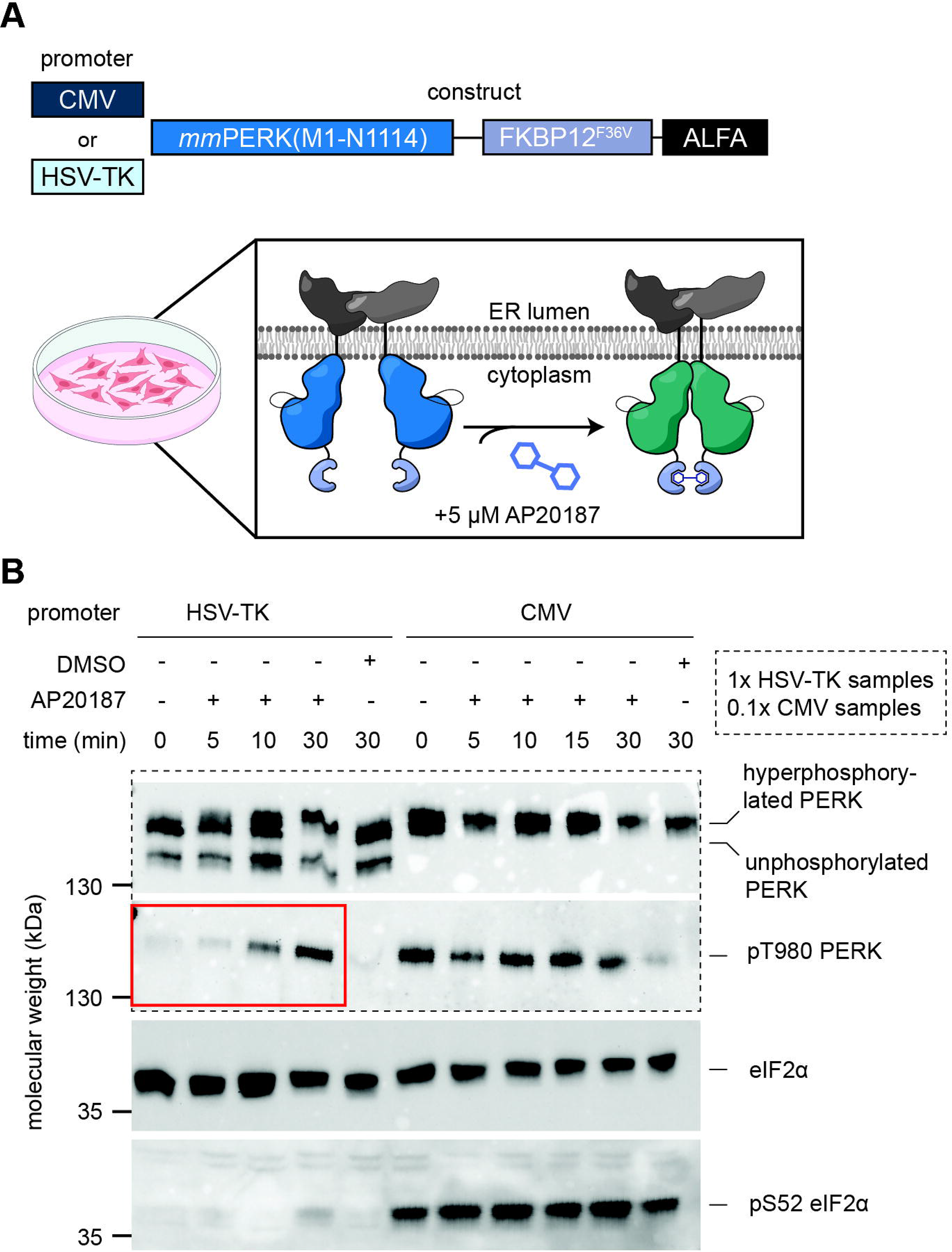
Dimerization is sufficient for PERK activation in cells. A. Schematic of constructs employed to test whether dimerization is sufficient for PERK activation in cells. Experimental strategy: HEK293 cells were transiently transfected with *Mm*PERK constructs and, 2 days post-transfection, dimerization was induced by addition of AP20187 (monomers: blue, dimers: green). B. Western blot of transiently-transfected HEK293 lysate, treated with dimerizer (AP20187) or vehicle control (DMSO) over a 30 min time course, stained for PERK, PERK pT980, eIF2α and eIF2α pS52. For lysates from the CMV condition 1/10 of the material was loaded to compensate for different expression levels. n = 2, representative image shown

To evaluate the effect of PERK dimerization on UPR activation, we employed western blot analysis to compare phosphorylation levels of the activation loop of PERK and eIF2α in dimerizer-treated (AP20187) and vehicle-treated (DMSO) cells. Under control of the CMV promoter, PERK migrates with a higher apparent molecular weight, consistent with previous reports of hyperphosphorylation upon PERK activation^1,48,59^ (Figure 3B). Strikingly, the addition of dimerizer to cells expressing PERK under the HSV-TK promoter induced a time-dependent increase in PERK activation loop autophosphorylation and low levels of eIF2α phosphorylation (Figure 3B, Supplementary Figure S7). In stark contrast, PERK overexpression under the CMV promoter led to both PERK and eIF2α phosphorylation that was independent of induced dimerization.

In summary, dimerization is sufficient to drive both PERK auto- and substrate (eIF2α) phosphorylation in cells.

### Substrate phosphorylation by PERK is dimerization-dependent

Having demonstrated that dimerization is necessary and sufficient for PERK autophosphorylation, we speculated that dimerization would also allosterically promote substrate phosphorylation. To deconvolute the effects of autophosphorylation from substrate phosphorylation, we first autophosphorylated both monomers (PERK^KD^-FKBP) and dimers (PERK^KD^-FKBP^AP^). Then, in a second step, we compared monomer and dimer-mediated eIF2α phosphorylation over time (Figure 4A).

**Figure 4.**
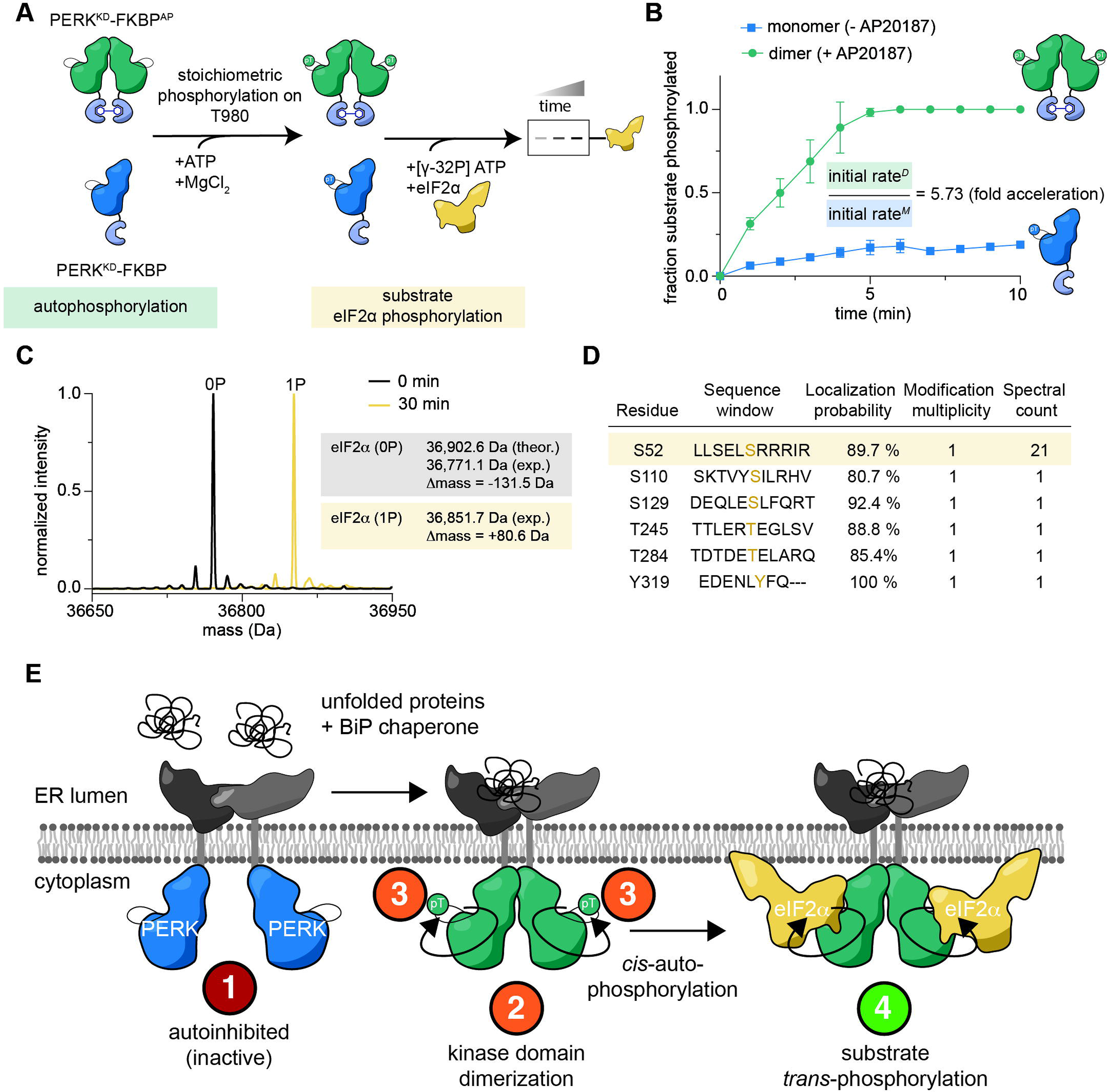
Substrate phosphorylation by PERK is dimerization-dependent. A. Experimental strategy to deconvolute substrate eIF2α phosphorylation from autophosphorylation. B. Time course of radiometric kinase assay of eIF2α phosphorylation by PERK^KD^-FKBP (monomeric, blue) and PERK^KD^-FKBP^AP^ (dimeric, green). C. Intact mass spectrometry of eIF2α after 30 min phosphorylation by PERK^KD^-FKBP^AP^. D. Tandem MS analysis of eIF2α to identify phosphosites after 30 min of phosphorylation by PERK^KD^-FKBP^AP^. E. Model of dimerization-stimulated *cis*-autophosphorylation of PERK. Unfolded proteins activate the UPR via the luminal domain in the ER, inducing conformational changes which trigger dimerization of the kinase domains on the cytoplasmic side of the ER. The back-to-back conformation is *cis*-autophosphorylation competent and autophosphorylates T980 in the activation loop. The dimeric, phosphorylated kinase is then primed for eIF2α binding and phosphorylation.

Dimerization of PERK increased the rate of substrate phosphorylation by approximately 6-fold compared to the monomeric kinase (Figure 4B), which is equivalent to the increase in the rate constant of autophosphorylation induced by dimerization (Figure 2G). We confirmed that eIF2α was exclusively and site-specifically phosphorylated on S52 (Figure 4C-D). This observation demonstrates that substrate phosphorylation, like autophosphorylation, is dimerization dependent.

Taking all of our findings together, we propose a model in which PERK is constitutively dimerized by its luminal domain, but its kinase domain is maintained in an inactive, monomeric conformation in the absence of proteotoxic stress (Figure 4E). The sensing of unfolded proteins in the ER lumen induces conformational changes that are transduced over the ER membrane to drive back-to-back dimerization and allosteric activation of the kinase domains. PERK then undergoes *cis*-autophosphorylation of both protomers, rendering it capable of binding and phosphorylating eIF2α.

### Dimerization-dependent *cis*-autophosphorylation activates the UPR and ISR kinases

The PERK kinase domain is evolutionarily most related to the ISR kinases HRI, Gcn2 and PKR^60^. As these kinases are structurally and functionally related, we hypothesized that they are all regulated by the same mechanism. Activation loop *cis*-autophosphorylation has already been demonstrated for PKR^38^, so we did not conduct any experiments on its kinase domain. Although the kinase domain of Ire1 is more distantly related, the crystal structure of the yeast Ire1 kinase domain exhibits a homologous back-to-back conformation that has been shown to be physiologically relevant for Ire1 signaling^37^. We therefore investigated whether HRI, Gcn2 and Ire1 also undergo dimerization-dependent *cis*-autophosphorylation using the same chemically induced dimerization system we have used to study PERK.

We used construct boundaries of a published crystal structure of *Saccharomyces cerevisiae* Gcn2^3^, in which the insertion loop between β4 and β5 in the N-lobe of the kinase domain was deleted (Gcn2^KD^, Figure 5A). Gcn2 heterodimers, purified according to the same scheme as PERK (Figure 1G), exhibited autophosphorylation only on the kinase-active protomer (Figure 5B). Intact mass spectrometry confirmed mono-phosphorylation exclusively of the active protomer (Figure 5C and Supplementary Table 2). Tandem mass spectrometry revealed phosphorylation of T882, corresponding to the canonical autophosphorylation site in its activation loop^32^ (Supplementary Figure 8A).

**Figure 5.**
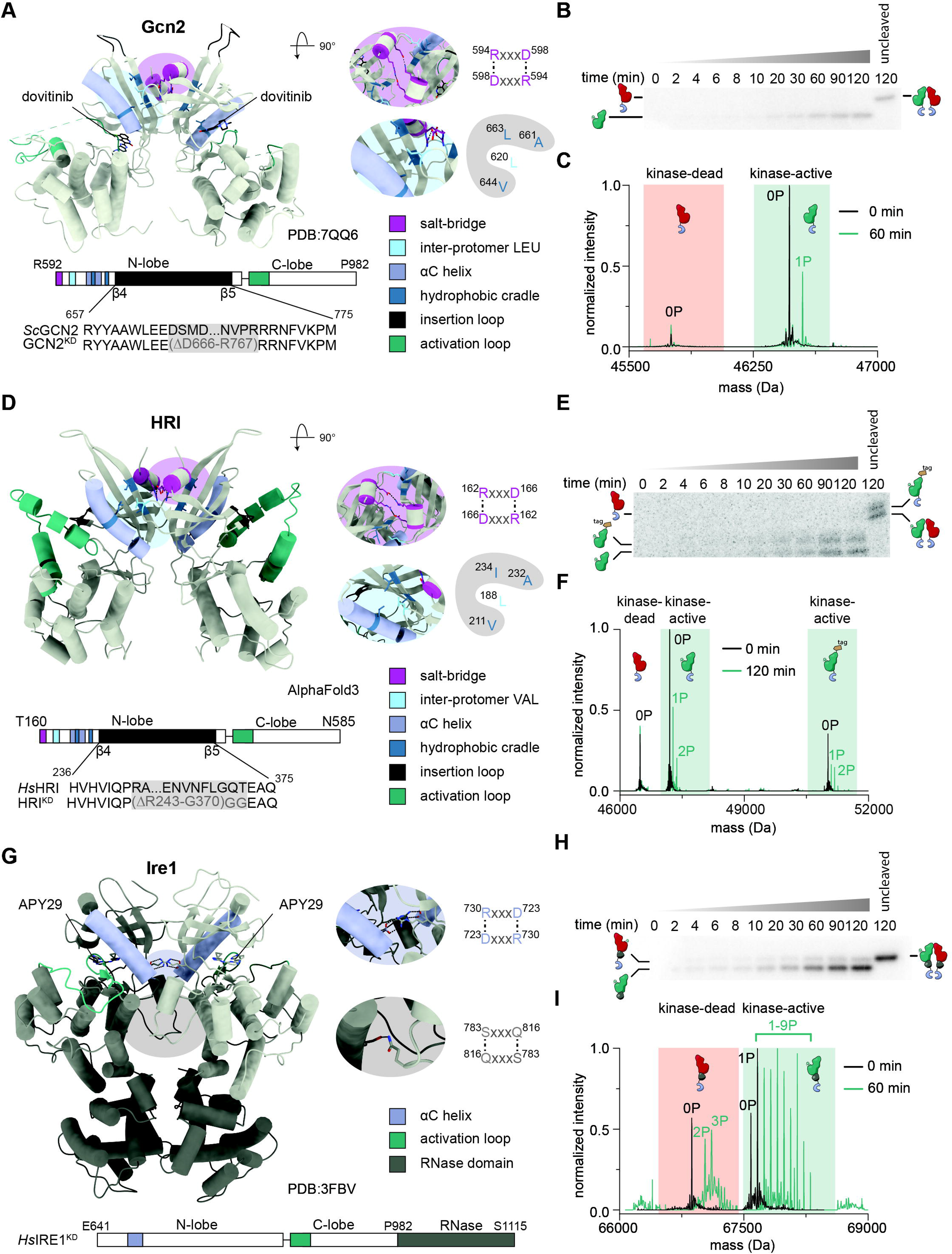
Dimerization-dependent *cis*-autophosphorylation is conserved in the UPR and ISR kinases. A. Crystal structure of dimeric *Sc*Gcn2 kinase domain in complex with dovitinib (PDB: 7QQ6). Relevant features for catalysis and dimerization in the kinase are highlighted in the structure and domain architecture (magenta: inter-dimer salt bridge, light blue: inter-protomer leucine (LEU), violet: α-C helix, dark blue: hydrophobic cradle, black: insertion loop in the kinase between β4 and β5, green: activation loop). With reference to the wildtype protein (*Sc*Gcn2), the deletion introduced in the insertion loop is depicted in the engineered Gcn2^KD^. B. Autophosphorylation time course of cleaved Gcn2 heterodimers. C. Intact mass spectrometry of Gcn2^KD^ heterodimers after 1 h autophosphorylation at 1 μM. D. AlphaFold prediction of dimeric *Hs*HRI^KD^. Relevant features for catalysis and dimerization in the kinase are highlighted in the structure and domain architecture (magenta: inter-dimer salt bridge, light blue: inter-protomer valine (VAL), violet: α-C helix, dark blue: hydrophobic cradle, black: insertion loop in the kinase between β4 and β5, green: activation loop). With reference to the wildtype protein (*Hs*HRI), the deletion introduced in the insertion loop is depicted in the engineered HRI^KD^. E. Autophosphorylation time course of cleaved HRI^KD^ heterodimers. The two bands correspond to partial cleavage of the kinase-active HRI protomer, which retains an affinity tag and its N-terminal methionine. F. Intact mass spectrometry of HRI^KD^ heterodimers before and after 2 h autophosphorylation at 10 μM. G. Crystal structure of dimeric *Sc*Ire1 kinase domain (PDB: 3FBV) in complex with APY29. Relevant features are highlighted (violet: α-C helix, green: activation loop, dark grey: RNase domain). Construct boundaries are depicted in the domain overview. H. Autophosphorylation time course of cleaved Ire1^KD^ heterodimers. I. Intact mass spectrometry of Ire1^KD^ heterodimers before and after 1 h autophosphorylation at 1 μM.

Lacking an experimental structure of HRI, we used AlphaFold^61^ to predict the structure of the HRI kinase domain dimer (Figure 5D). The obtained model revealed a back-to-back conformation superimposable with experimental structures of PERK, PKR and Gcn2. As for PERK and Gcn2, we modified the ISR kinase-specific insertion loop to remove off-target phosphorylation sites. Like PERK and Gcn2, heterodimers of active and inactive HRI exhibited autophosphorylation only on the kinase-active protomer (Figure 5E). The double band visible in the autoradiograph resulted from incomplete cleavage of the affinity tag in the active kinase protomer (Figure 5E). The identity of all species and their phosphorylation signatures were further confirmed by intact and tandem mass spectrometry (Figure 5F and Supplementary Table 2). The catalytically dead protomer remained unphosphorylated, whereas the catalytically active protomer was modified with up to two phosphates in the activation loop (Supplementary Figure 8B).

Finally, we investigated Ire1, which is further distinguished from the ISR kinases by the presence of a RNase domain C-terminal to the kinase domain (Figure 5G). We employed the same construct boundaries as Korennykh et al^37^. Ire1^KD-RNase^ heterodimers exhibited two bands post-autophosphorylation: a high intensity band for the catalytically active subunit, and a faint band for the catalytically inactive subunit, with autophosphorylation of both increasing over time (Figure 5H). Mass spectrometry confirmed the incorporation of up to 9 phosphates on the catalytically active protomer and 2-3 phosphates on the catalytically inactive protein (Figure 5I, Supplementary Figure 8C, and 8D and Supplementary Table 2). Unlike the ISR kinases, Ire1 exhibited limited *trans*-autophosphorylation activity over time. Nevertheless, the marked preference for phosphorylation of the catalytically active protomer indicates that *cis*-autophosphorylation is the dominant reaction.

In summary, the UPR and ISR kinases are all activated by dimerization-dependent *cis*-autophosphorylation.

## Discussion

In this study, we show that the UPR and ISR kinases PERK, Gcn2, HRI and Ire1 are activated by intramolecular *cis*-autophosphorylation of their activation loops. We have determined that *cis*-autophosphorylation of PERK is dimerization-dependent and identify specific residues in the back-to-back dimer interface that allosterically drive kinase activity. We rationalize the mechanism of dimerization-induced *cis*-autophosphorylation in a quantitative model, benchmarking it with kinetic parameters. Furthermore, we show that inducible dimerization of PERK is necessary and sufficient to activate the UPR in cells. Finally, we demonstrate that dimerization-stimulated *cis*-autophosphorylation is conserved across all of the UPR and ISR kinases.

Our findings imply that dimers are the functional signaling units in the UPR and ISR. To date, PERK and Ire1 have been proposed to rely on oligomerization and clustering for their activation by *trans*-autophosphorylation. To achieve acute and high-fidelity signal transduction, such a mechanism would require the rapid translocation of transmembrane proteins in a crowded membrane in order to assemble and disassemble active clusters in response to signaling cues. Furthermore, the resolution of face-to-face dimers following *trans*-autophosphorylation would be required in order to permit substrate phosphorylation. Back-to-back dimers that *cis*-autophosphorylate obviate the requirement to form face-to-face dimers, thereby solving this problem. With respect to the active conformation, multiple reports have highlighted the importance of the back-to-back dimer in the UPR and ISR kinases. For Ire1, it has been suggested that dimerization of the luminal domain is necessary and sufficient for its activation^41^. As opposed to cluster formation of over-expressed Ire1, a quantitative imaging study of endogenous Ire1 is notable for the absence of higher-order clusters, despite Ire1 autophosphorylation and UPR activation^62^. For the ISR kinases, formation of the conserved inter-protomer salt-bridge on the N-lobe has previously been implicated in autophosphorylation and activation of the kinases^63^. Recently, it was discovered that sub-stoichiometric quantities of ATP-competitive PERK inhibitors paradoxically activate Gcn2^64^, which can be explained by the bidirectional allosteric crosstalk between kinase protomers. Occupancy of one protomer of the dimer by an ATP-competitive drug with typically higher affinity than ATP shifts the kinase conformation towards the active conformation, thereby promoting dimerization-dependent allosteric activation of the opposing protomer. Finally, dimerization-dependent *cis*-autophosphorylation of PKR has already been described in yeast^38^. In summary, activation loop *cis*-autophosphorylation provides a unifying framework that reconciles structural, biochemical and cellular observations across the UPR and ISR.

*Cis*-autophosphorylation, however, implies the requirement for a mechanism to maintain these kinases in an inactive conformation. Kinases can be kept inactive in numerous ways, including inhibition by another protein, autoinhibition by intrinsic regulatory domains, or stabilization of an inactive conformation of the activation loop itself. For the UPR and ISR kinases, a discrete inactive conformation of the activation loop has, to date, not been described. However, given the precedent for inactive conformations of the activation loop in other kinases^65,66^, it is eminently plausible that such a mechanism inactivates the ISR and UPR kinases. Presumably, dimerization of the kinase domains and the associated conformational changes in the αC helix that promote activation would be coupled to destabilization of the inactive conformation of the activation loop. Further work will undoubtedly be required to elucidate the mechanism by which these kinases are inactivated in the absence of stress. It is worth noting that, although a phosphatase has been identified for eIF2α^67–69^, none has been reported to dephosphorylate the activation loop of any UPR or ISR kinase.

Dimerization and oligomerization in the context of the crowded cellular environment are equally affected by concentration and barriers to diffusion. Therefore, the advantages of a dimerization-dependent *cis*-autophosphorylation mechanism over the oligomerization-mediated *trans*-phosphorylation mechanism proposed by others only holds true if the UPR and ISR kinases assemble as inactive dimers in the cell. It is therefore notable that Ire1 has been suggested to form preformed dimers^62^, Gcn2 has been reported to be constitutively dimeric, adopting an anti-parallel, dimeric inactive fold under homeostatic conditions^26^, PKR has been shown to dimerize via its RNA-binding domain^70,71^, and HRI has been shown to be predominantly dimeric in solution^39^. Homologous dimeric assemblies have previously been described for the bacterial kinase PknB^72^ and the mitotic spindle kinase Nek7^73,74^. Although Nek7 lacks an intrinsic dimerization domain, its back-to-back dimerization and activation is facilitated by Nek9. These observations suggest that constitutive dimerization mediated via diverse sensory domains creates a pre-organized platform necessary and sufficient for signal transduction.

While the mechanism of allosteric activation of the UPR in the lumen of the ER remains elusive, the mechanism we propose for PERK and Ire1 activation shifts the focus from oligomerization and clustering to allosteric regulation. Although activated by distinct physiological cues, the UPR and ISR kinases share a conserved molecular activation logic (Figure 6). We propose that, in the absence of stress, the kinases adopt an inactive conformation in which the activation loop occludes the catalytic cleft and the αC helix is displaced from its active conformation, precluding substrate binding and catalysis (Figure 6, inactive, red). Back-to-back dimerization of the kinase domain, elicited by conformational changes in the sensor domain(s), establishes conserved inter-protomer contacts which allosterically expel the activation loop and position the αC helix in an active conformation compatible with *cis*-autophosphorylation (dimerization-dependent *cis*-autophosphorylation, orange). Activation loop autophosphorylation stabilizes the αD helix in a substrate binding-competent conformation (active, green), thereby driving dimerization-dependent substrate phosphorylation (green).

**Figure 6.**
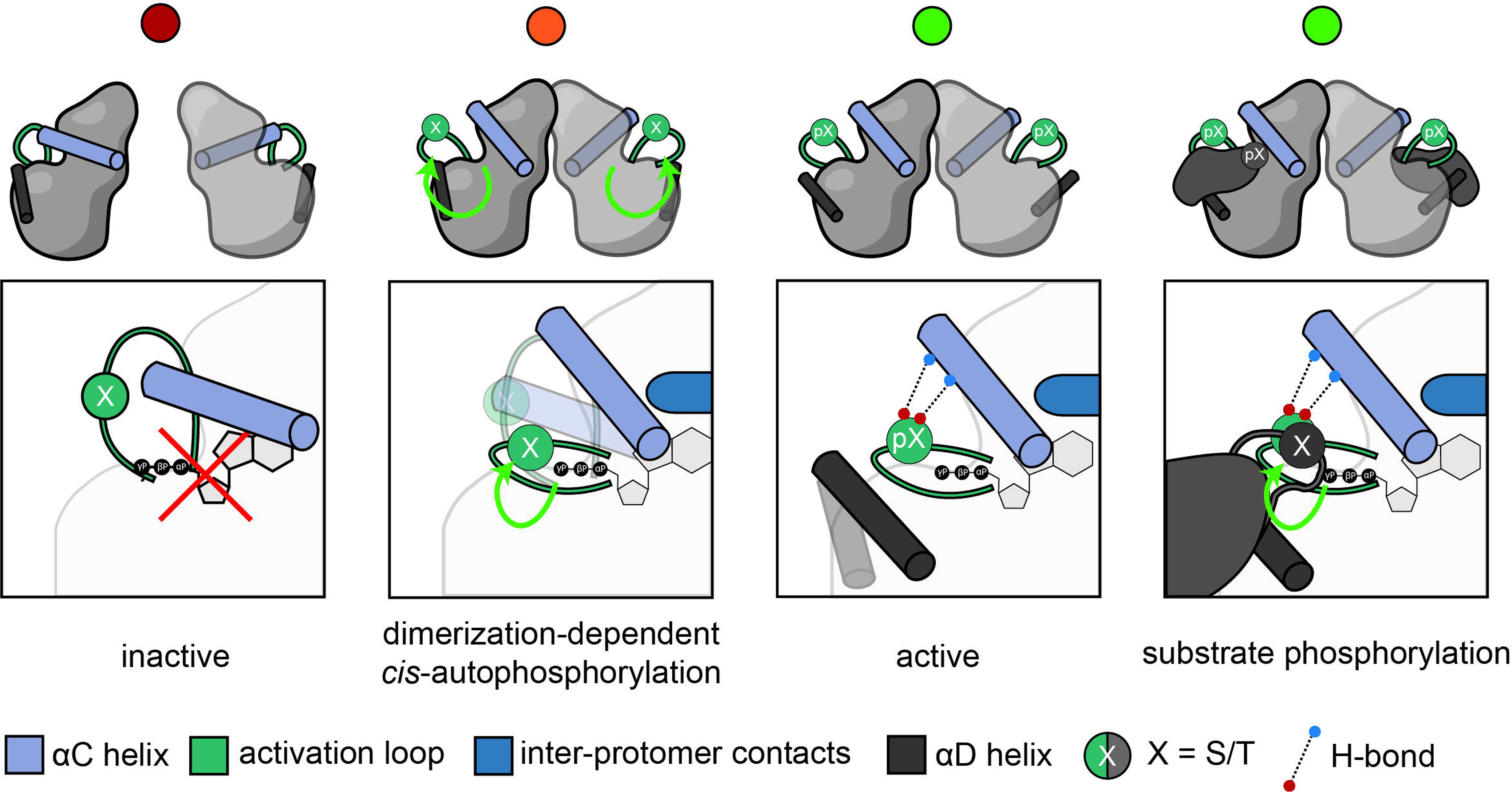
Model of back-to-back *trans*-activated *cis*-autophosphorylation. In the absence of proteotoxic stress, protein kinases adopt an inactive, monomeric conformation in which the conformations of the activation loop (dark green) and αC helix (purple) preclude autophosphorylation. Back-to-back dimerization (blue) allosterically triggers displacement of the activation loop and repositioning of the αC helix into a catalytically competent conformation, which permits *cis*-autophosphorylation of the activation loop S/T residue(s). Activation loop autophosphorylation stabilizes the active conformation of the kinase and promotes binding and subsequent phosphorylation of its substrate (grey).

Open questions that require further investigation revolve around the allosteric changes in the luminal domains of PERK and Ire1 upon proteotoxic stress, including the role of BiP and other ER chaperones, and how they elicit dimerization of the cytoplasmic kinase domains via the transmembrane helix. Previous studies suggest that changes in lipid composition can also activate UPR signaling by conformational rearrangement of the transmembrane helices^75,76^, a phenomenon which implies that local allosteric changes are sufficient to drive kinase activation. Similarly, for the non-transmembrane ISR kinases PKR, Gcn2, and HRI, work remains to elucidate the mechanisms by which the sensing of their input signals results in the concerted conformational changes required for kinase activation.

In summary, our work resolves important questions regarding the activation of four important stress response kinases, whose dysregulation is associated with human disease. The mechanistic framework we have established is likely a blueprint for the activation of other kinases not investigated in this study.

## Materials and Methods

### Radiometric kinase assays

Autophosphorylation and substrate phosphorylation assays were performed using radiolabeled [γ-32P] ATP (Hartman Analytic). All reactions, except for Ire1 autophosphorylation, were performed in the same kinase reaction buffer: 150 mM NaCl, 50 mM Tris pH 7.4, 1 mM TCEP, 0.25% CHAPS, whereas the buffer composition for Ire1 was 650 mM NaCl, 50 mM Tris pH 7.4, 1 mM TCEP, 0.25% CHAPS. Autophosphorylation assays were started by adding ATP and MgCl_2_, spiked with 1.6 μL of [γ-32P] ATP per 100 μL final reaction volume, resulting in a final reaction mix composition of 150/650 mM NaCl, 50 mM Tris pH 7.4, 1 mM TCEP, 0.25% CHAPS, 1 mM ATP, 2 mM MgCl_2_. All assays except for substrate eIF2α phosphorylation were incubated at 23°C in a thermoblock, with shaking at 300 rpm. Samples for eIF2α substrate phosphorylation assays were prepared as follows: stoichiometric autophosphorylation of monomeric PERK kinase (PERK^KD^-FKBP) was performed in kinase reaction buffer for 3 h at 23°C at 10 μM, whereas the dimeric kinase (PERK^KD^-FKBP^AP^) was autophosphorylated for 20 minutes at 23°C at 1 μM. Autophosphorylated PERK^KD^-FKBP(AP) at 1 μM was then mixed with eIF2α spiked with 1.6 μL of [γ-32P] ATP per 100 μL final reaction volume to initiate substrate phosphorylation on ice. For all assays, each time point was sampled in 2 μL of 0.5M EDTA to quench the reaction. For all heterodimer autophosphorylation assays, 3 μL of 1 mg/mL 3C protease was added per 12 μL quenched sample for 3 h at room temperature.

Samples of autophosphorylated PERK were analyzed by SDS-PAGE as well as spotting on 0.45 μm nitrocellulose membrane (Cytiva), whereas all other assays were analyzed by SDS-PAGE. For SDS-PAGE, samples were mixed with 3 μL of 5x loading dye and separated with 12% SDS-PAGE gels. Gels were run for 35 minutes at 150V, before washing them 3×5 min in H_2_O before drying at 80°C for 30 minutes under vacuum. After membrane spotting, the nitrocellulose membrane was washed 3x 5min in 75 mM H_3_PO_4_. Both gels and membranes were then wrapped in Saran wrap and exposed on a phosphoscreen overnight. The screen was imaged with an Amersham Typhoon and the radioactive signal was quantified in ImageJ.

### Isothermal titration calorimetry

ITC measurements were performed on a Microcal PEAQ-ITC (Malvern) at 25°C with constant stirring. PERK^KD^^30^ was concentrated to 300 – 400 μM and degassed for 10 minutes. PERK^KD^ concentration was measured on a spectrofluorimeter at 280 nm and aggregation was monitored at 333 nm in triplicates before calculating the final concentration. PERK^KD^ was titrated into buffer and heat of dilution was subtracted from the raw thermogram. Data was globally fitted with Affinimeter’s dissociation model [Free Species] A_2_ ^77,78^. Data points fitted and excluded for analysis are highlighted in Supplementary Figure 6.

### Cell culture and transfection

HEK293 cells were cultured in DMEM supplemented with 5% FBS and ZellShield following the manufacturer’s instructions (Minerva Biolabs). PERK^FL^-3C-FKBP12^F36V^-TEV-ALFA constructs were cloned in either the pBiT1.1-C [TK/LgBiT]^79^ (HSV-TK) or the pCMV backbone and transfected using Lipofectamine 3000 (Invitrogen). Cell medium was exchanged 24 h post-transfection. Cells were harvested in RIPA buffer supplemented with phosphatase inhibitors (150 mM NaCl, 10 mM Tris pH 7.4, 0.1% SDS, 0.1% Triton X100, 0.1% w/v sodium deoxycholate, 1x protease inhibitor cocktail (Sigma P8849), 10 mM beta glycerophosphate, 2 mM sodium pyrophosphate, 1 mM sodium orthovanadate, 50 mM sodium fluoride) and lysed by three freeze/thaw cycles. Cells were sonicated for 2 sec on ice and total protein of whole cell lysate was measured using the Qubit™ Protein broad range kit (Thermo Fisher Scientific). 3 μg (CMV samples) or 30 μg (HSV-TK samples) total protein was loaded per lane on 8% SDS-PAGE gels for blotting against PERK and pT980 PERK. 30 μg (CMV and HSV-TK) were loaded on 12% SDS-PAGE gels for blotting against eIF2α and pS52 eIF2α. Gels were blotted overnight at 150 mA, blocked for 2 h in 5% milk powder in TBS-T (TBS + 0.05% Tween), and primary antibodies (PERK: rabbit anti-PERK, Cell Signaling technologies, Cat. #3192, 1:1000; pT980 PERK: rabbit anti-phospho PERK, Cell Signaling Technologies, Cat. #3179, 1:1000, eIF2α: rabbit anti-eIF2α, Cell Signaling Technologies, Cat. #9722, 1:1000; pS52 eIF2α: rabbit anti-phospho eIF2α, cell Signaling Technologies, Cat. #9721, 1:1000) were incubated overnight at 4°C with constant rotation. Blots were washed 3x 5 min in TBS-T, secondary antibodies (goat anti-rabbit IgG (H+L), HRP coupled, invitrogen, Cat. #31460, 1:5000) were incubated for 2 h at room temperature before washing again 3x 5 min. Western blots were developed with ECL Western blotting substrate (Promega W1001) and imaged using a ChemiDoc (BioRad).

## Acknowledgements

Proteomics analyses were performed on instruments of the Vienna BioCenter Core Facilities (VBCF). ITC, nDSF and CD experiments were performed on instruments of the Protein Technologies Facility of the Vienna BioCenter Core Facilities (VBCF). We thank Affinimeter for granting a temporary license for ITC data analysis. Cell culture medium was prepared by the Max Perutz labs media kitchen. We thank David Drescher and Dr. Stefan Schüchner of the Ogris lab for their valuable input on antibodies, Dr. Ingrid Frohner for *Saccharomyces cerevisiae* gDNA, and Dr. Jonas Ries for input on the kinetic modeling. The FKBP12^F36V^ plasmid was a kind gift of the Matos lab (Max Perutz labs). We acknowledge members of the Leonard lab for constructive feedback and input on the manuscript. This work was funded by the Austrian Science Fund (FWF) grants P36212, and P36724 to T.A.L and a Max Perutz PhD Fellowship to M.O.

## Author contributions

M.O. performed all experimental work reported in this manuscript. T.L. and M.O. derived the kinetic model of *trans*-activated *cis*-autophosphorylation. M.O. and T.L. wrote and edited the manuscript. M.O. and T.L. obtained funding for the study.

## Resource availability

### Correspondence

Further information and requests for resources and reagents should be directed to and will be fulfilled by corresponding author, Thomas A. Leonard.

### Data availability

The mass spectrometry proteomics data have been deposited to the ProteomeXchange Consortium via the PRIDE^80^ partner repository with the dataset identifiers PXD081199, PXD081308, PXD081379, PXD081533, and PXD081379.

This paper does not report original code.

Any additional information required to reanalyse the data reported in this paper is available from the corresponding author upon request.

All data are available in the main text or the supplemental information.

