## Supplementary Information for "Dimerization-dependent *cis*-autophosphorylation activates the UPR and ISR kinases"

Contents:

Supplementary Figures 1-8

Supplementary Tables 1-2

Supplementary Methods

Supplementary References

Figure S1. PERK construct optimization for stoichiometric activation loop phosphorylation.

**A**

PERK$^{KD}$-FKBP$^{AP}$ (H545-G1082, ΔT701-F865)
(pre-autophosphorylation)

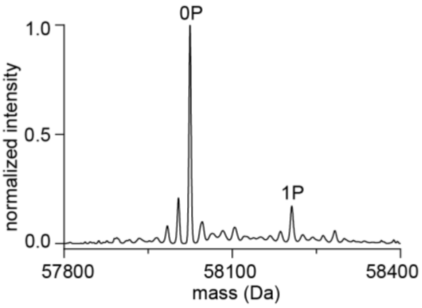

**B**

PERK$^{KD}$-FKBP$^{AP}$ (H545-G1082, ΔT701-F865)
(post-autophosphorylation)

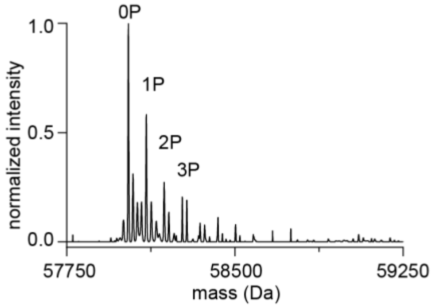

**C**

| Residue | Sequence window | Localization probability | Modification multiplicity | Spectral count |
|---|---|---|---|---|
| T553 | RKESETQCQTE | 94.8 % | 1;2;3 | 40 |
| T557 | ETQCQTESKYD | 95.0 % | 1;2;3 | 45 |
| S700 | RMDPFSTKNTV | 95.8 % | 1;2;3 | 130 |
| T701 | MDPFSTKNTVG | 95.8 % | 1;2;3 | 177 |
| T869 | FSTKNTVGQLQ | 95.8 % | 1;2;3 | 149 |
| T980 | AYATHTGQVGT | 94.7 % | 1;2 | 72 |

(spectral counts below 40 not shown)

**D**

PERK$^{KD}$-FKBP$^{AP}$ (G580-G1082, ΔS700-T869)
(pre-autophosphorylation)

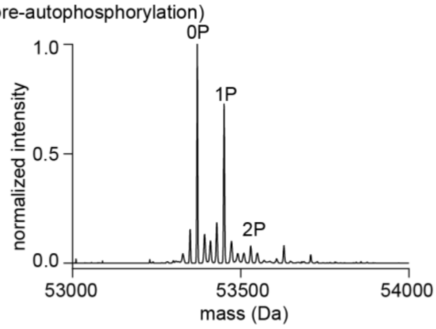

**E**

PERK$^{KD}$-FKBP$^{AP}$ (G580-G1082, ΔS700-T869)
(co-expression with lambda phosphatase) (pre-autophosphorylation)

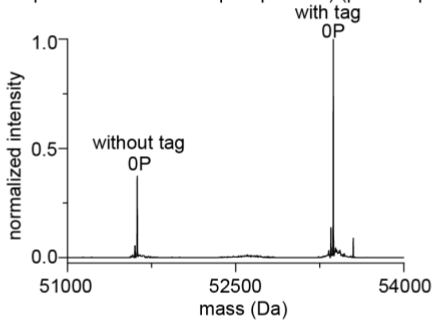

**F**

PERK$^{KD}$-FKBP$^{AP}$ (G580-G1082, ΔS700-T869)
(post-autophosphorylation)

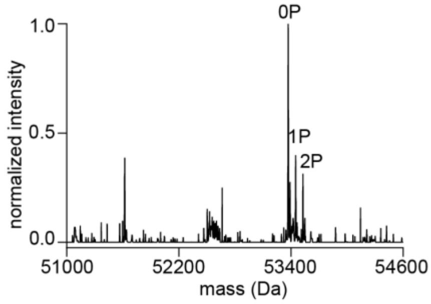

**G**

PERK$^{KD}$-FKBP$^{AP}$ (G580-G1082, ΔS700-T869, S681G, S682G, S684G, S690G) (pre-autophosphorylation)

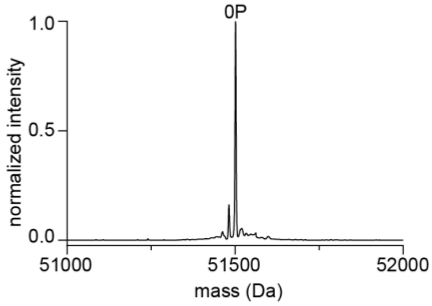

**H**

PERK$^{KD}$-FKBP$^{AP}$ (G580-G1082, ΔS700-T869, S681G, S682G, S684G, S690G) (post-autophosphorylation)

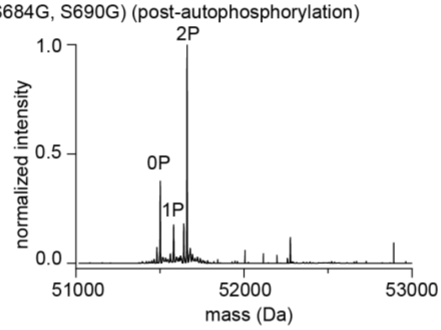

**I**

| Residue | Sequence window | Localization probability | Modification multiplicity | Spectral count |
|---|---|---|---|---|
| T676 | LKDESTDWPLG | 100 % | 1;2 | 13 |
| T944 | SNIFFTMDDVV | 94.6 % | 1 | 5 |
| T968 | DEEEQTVLTPM | 94.3 % | 1;2 | 9 |
| Y976 | TPMPAYATHTG | 90.1 % | 1;2 | 28 |
| T978 | MPAYATHTGQV | 92.2 % | 2 | 15 |
| T980 | AYATHTGQVGT | 93.6 % | 1;2 | 125 |

(spectral counts below 5 not shown)

**J**

PERK$^{KD}$-FKBP$^{AP}$ (G580-G1082, ΔS700-T869, S675G, S681G, S682G, S684G, S690G) (pre-autophosphorylation)

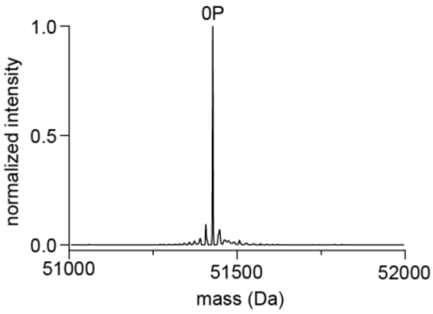

A. Intact mass spectrometry of dimeric PERK$^{KD}$-FKBP$^{AP}$ (H545-G1082, ΔT701-F865).
B. Intact mass spectrometry of autophosphorylated, dimeric PERK$^{KD}$-FKBP$^{AP}$ (H545-G1082, ΔT701-F865).
C. Tandem mass spectrometry analysis of autophosphorylated, dimeric PERK$^{KD}$-FKBP$^{AP}$ (H545-G1082, ΔT701-F865).
D. Intact mass spectrometry of dimeric PERK$^{KD}$-FKBP$^{AP}$ (G580-G1082, ΔS700-T869).
E. Intact mass spectrometry of dimeric PERK$^{KD}$-FKBP$^{AP}$ (G580-G1082, ΔS700-T869), purified after co-expression with lambda phosphatase.
F. Intact mass spectrometry of autophosphorylated, dimeric PERK$^{KD}$-FKBP$^{AP}$ (G580-G1082, ΔS700-T869).
G. Intact mass spectrometry of dimeric PERK$^{KD}$-FKBP$^{AP}$ (G580-G1082, ΔS700-T869, S681G, S682G, S684G, S690G).
H. Intact mass spectrometry of autophosphorylated, dimeric PERK$^{KD}$-FKBP$^{AP}$ (G580-G1082, ΔS700-T869, S681G, S682G, S684G, S690G).
I. Tandem mass spectrometry analysis of autophosphorylated, dimeric PERK$^{KD}$-FKBP$^{AP}$ (G580-G1082, ΔS700-T869, S681G, S682G, S684G, S690G).
J. Intact mass spectrometry of dimeric PERK$^{KD}$-FKBP$^{AP}$ (G580-G1082, ΔS700-T869, S675G, S681G, S682G, S684G, S690G).

#### Figure S2. Mass photometry of PERK$^{KD}$-FKBP$^{AP}$.

**A**

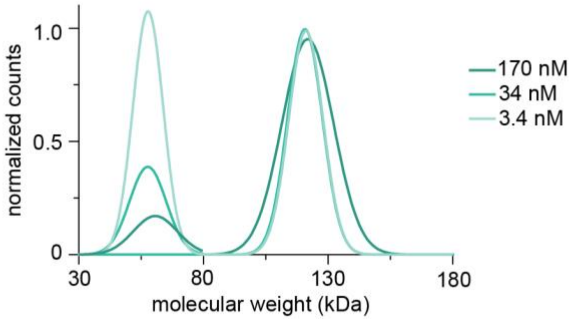

A. Gaussian fit of mass photometry of PERK$^{KD}$-FKBP$^{AP}$ (G580-G1082, ΔS700-T869, S675G, S681G, S682G, S684G, S690G).

#### Figure S3. Mass spectrometry of mutant PERK$^{KD}$ constructs employed in this study.

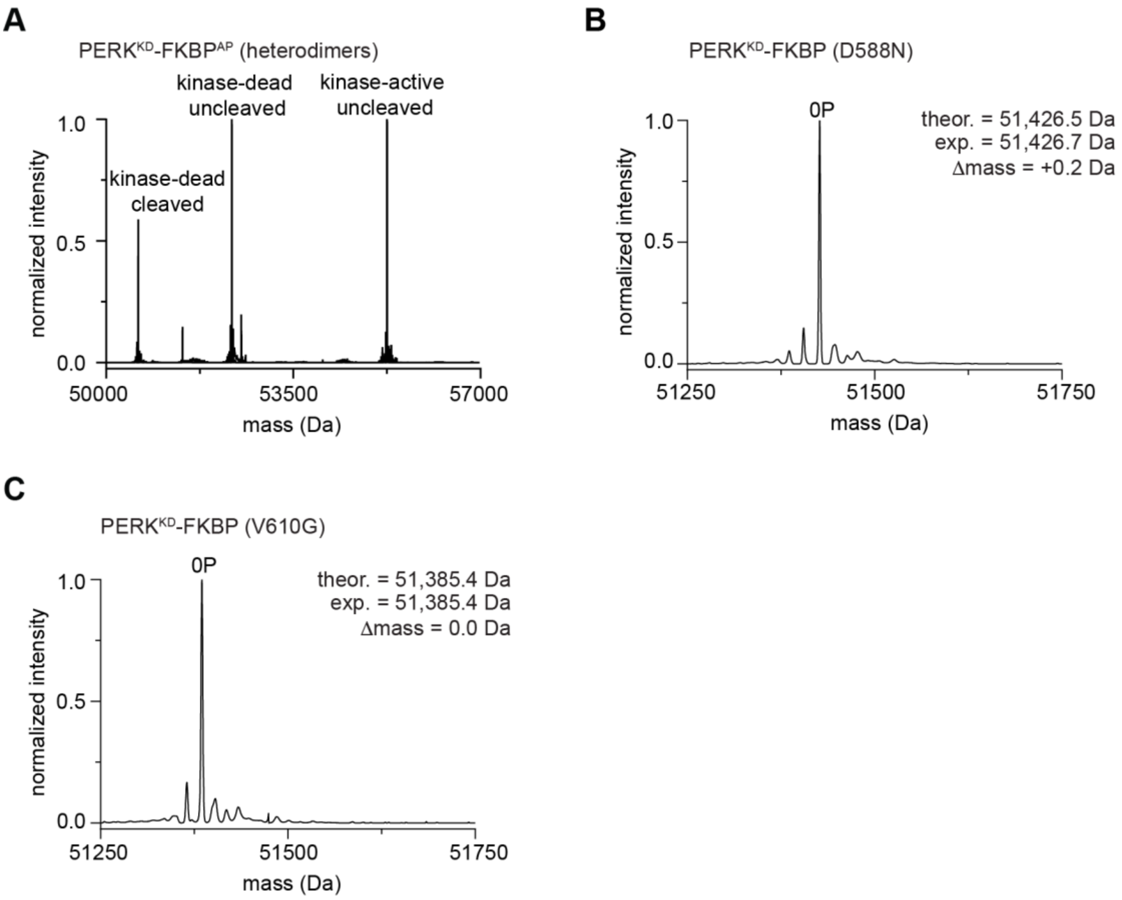

A. Intact mass spectrometry of PERK$^{KD}$-FKBP$^{AP}$ heterodimers comprised of kinase-active and kinase-dead protomers.
B. Intact mass spectrometry of dimeric PERK$^{KD}$-FKBP D588N.
C. Intact mass spectrometry of dimeric PERK$^{KD}$-FKBP V610G.

#### Figure S4. Biophysical characterization of PERK$^{KD}$-FKBP dimer interface mutants.

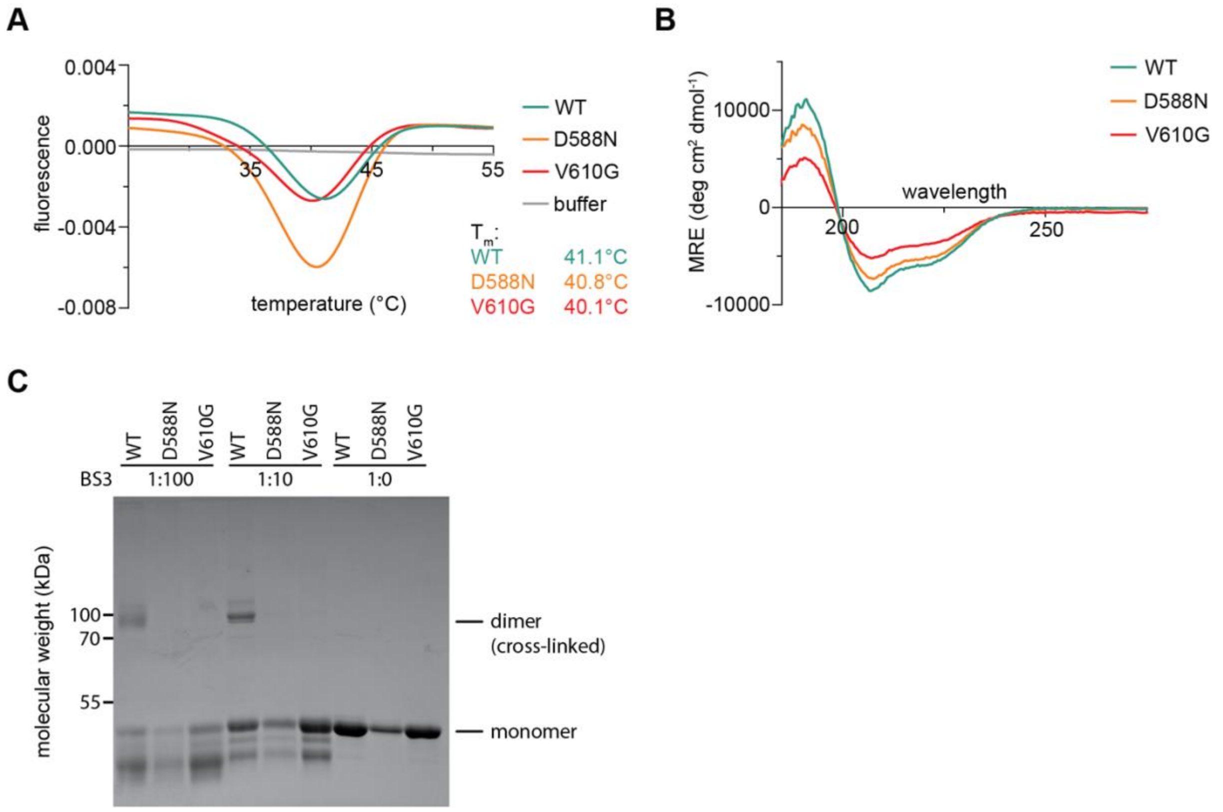

A. Nano differential scanning fluorimetry of apo, monomeric wildtype PERK$^{KD}$-FKBP, PERK$^{KD}$-FKBP D588N and PERK$^{KD}$-FKBP V610G.
B. Circular dichroism spectra of apo, monomeric wildtype PERK$^{KD}$-FKBP, PERK$^{KD}$-FKBP D588N and PERK$^{KD}$-FKBP V610G.
C. SDS-PAGE analysis of BS3 crosslinked wildtype, monomeric PERK$^{KD}$-FKBP, PERK$^{KD}$-FKBP D588N and PERK$^{KD}$-FKBP V610G in the presence of ATPγS. 1:100, 1:10, 1:0 = BS3:protein stoichiometry.

#### Figure S5. Kinetic model of dimerization-dependent *cis*-autophosphorylation.

**A**

**Definition and units of all variables:**

| | | |
|---|---|---|
| $[M]$ | concentration monomers | mol/L |
| $[D]$ | concentration dimers | mol/L |
| $K_D$ | affinity of homodimerization | mol/L |
| $k_{cat}$ | catalytic rate constant | $1/s^{-1}$ |
| $k_{obs}$ | experimentally observed pseudo-first order rate constant of autophosphorylation | $1/s^{-1}$ |
| $f_P$ | fraction of phosphorylated molecules | 0-1 |
| $f_D$ | fraction of dimers | 0-1 |
| $t$ | time | s |

**B**

##### Affinity of homodimerization

Phosphorylation does not affect the **affinity of dimerization** and the **rate constant, $k_{cat}$, of phosphorylation is the same for homodimer and heterodimers**, therefore monomer (M) and dimer (D) are always in equilibrium and do not change over time:

$$M + M \rightleftharpoons D$$

$$K_D = \frac{[M]^2}{[D]} \quad \text{rearrange} \quad [D] = \frac{[M]^2}{K_D} \qquad (1)$$

For weak dimerization, the fraction of molecules that are in dimers, $f_D$, is:

$$f_D \cong \frac{[2D]}{[M]} \qquad (2)$$

##### Catalytic rate constant ($k_{obs}$)

Only molecules that are in dimers can autophosphorylate. Each dimer contains 2 molecules, so the concentration of phosphorylation-competent molecules is:

$$[2D] = \frac{2[M]^2}{K_D} \qquad (3)$$

Substituting (3) into (2):

$$f_D = \frac{2[M]^2/KD}{[M]} = \frac{2[M]}{K_D} \qquad (4)$$

Rate of phosphorylation is given by the phosphorylation-competent molecules in the dimer:

$$k_{obs} = f_D k_{cat} = \frac{2[M]k_{cat}}{K_D} \qquad (5)$$

##### Kinase autophosphorylation over time

For a simple first order reaction to completion, the fraction of molecules phosphorylated, $f_P$, accumulates according to:

$$\frac{df_P}{dt} = k_{obs}(1 - f_P) \qquad (6)$$

Solving:

separate variables $\quad \frac{df_P}{1 - f_P} = k_{obs}dt$

integrate $\quad \int \frac{df_P}{1 - f_P} = \int k_{obs}dt$

$$-\ln(1 - f_P) = k_{obs}t + C$$

initial condition (no phosphorylation at time 0) $\quad f_P(0) = 0$

$$-\ln(1 - 0) = C$$

so $\quad C = 0$

therefore $\quad -\ln(1 - f_P) = k_{obs}t$

multiply by -1 $\quad \ln(1 - f_P) = -k_{obs}t$

exponentiate $\quad 1 - f_P = e^{-k_{obs}t}$

rearrange $\quad f_P(t) = 1 - e^{-k_{obs}t} \qquad (7)$

Substituting (5) into (7):

$$f_P(t) = 1 - e^{-\left(\frac{2*[M]*k_{cat}}{K_D}\right)t} \qquad (8)$$

A. List of all parameters defined in the model with respective units.
B. Full derivation of a mathematical model for dimerization-dependent *cis*-autophosphorylation.

#### Figure S6. Kinetic model of dimerization-dependent *cis*-autophosphorylation.

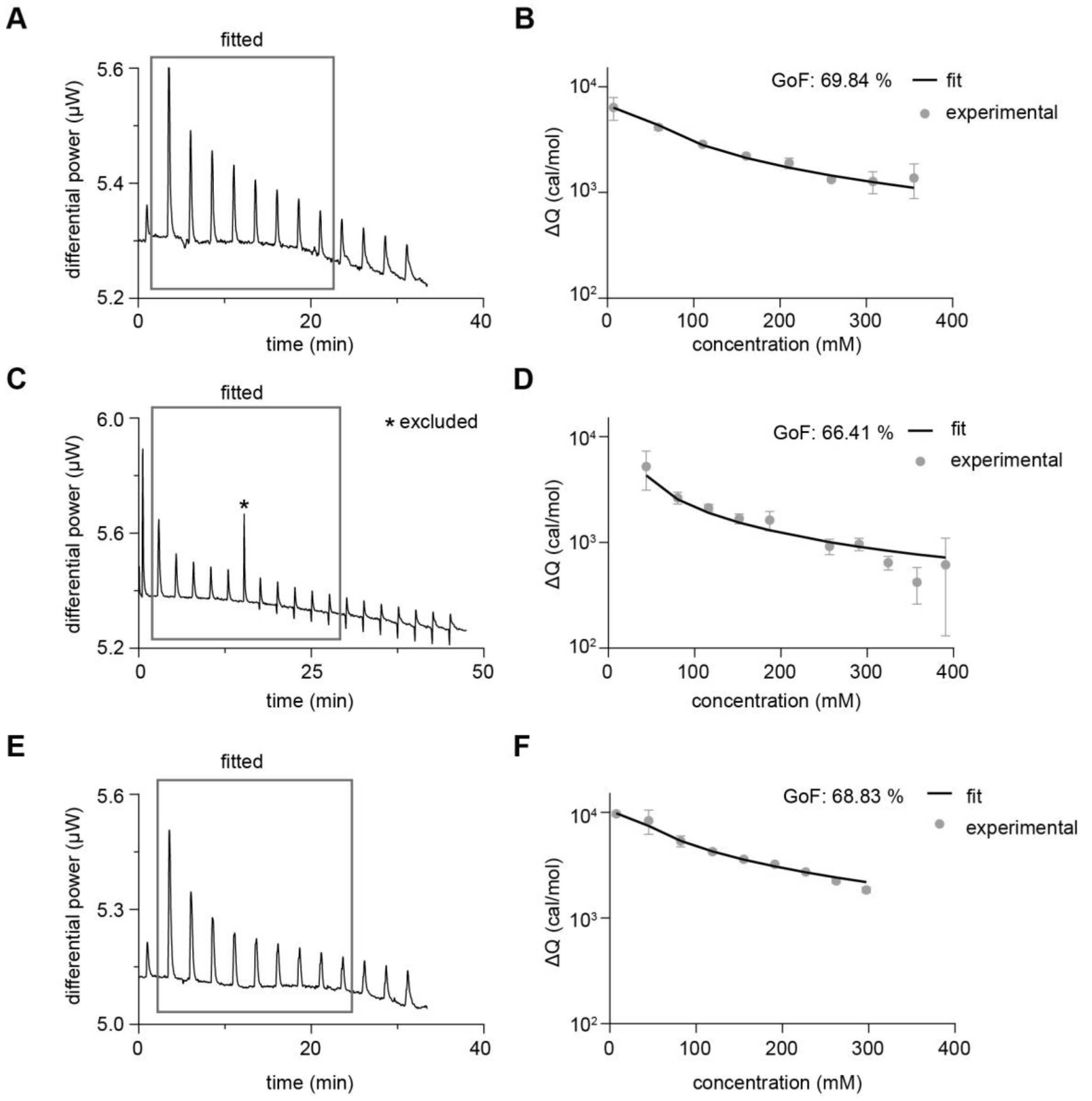

A. Raw ITC data normalized to baseline (buffer). Injections used for fitting are highlighted in a grey box and excluded datapoints are marked with a star (*).
B. Global fit to integrated heat upon injection (dots) with respective error bars and GlobalFit parameter. Fitting was done using Affinimeter's one-sided dissociation model[1]. The dissociation constant for homodimerization was determined from the fit.
C. Raw ITC trace for replicate 2.
D. Global fit to raw data of replicate 2.

E. Raw ITC trace for replicate 3.
F. Global fit to raw data of replicate 3.

#### Figure S7. PERK undergoes activation loop *cis*-autophosphorylation in cells.

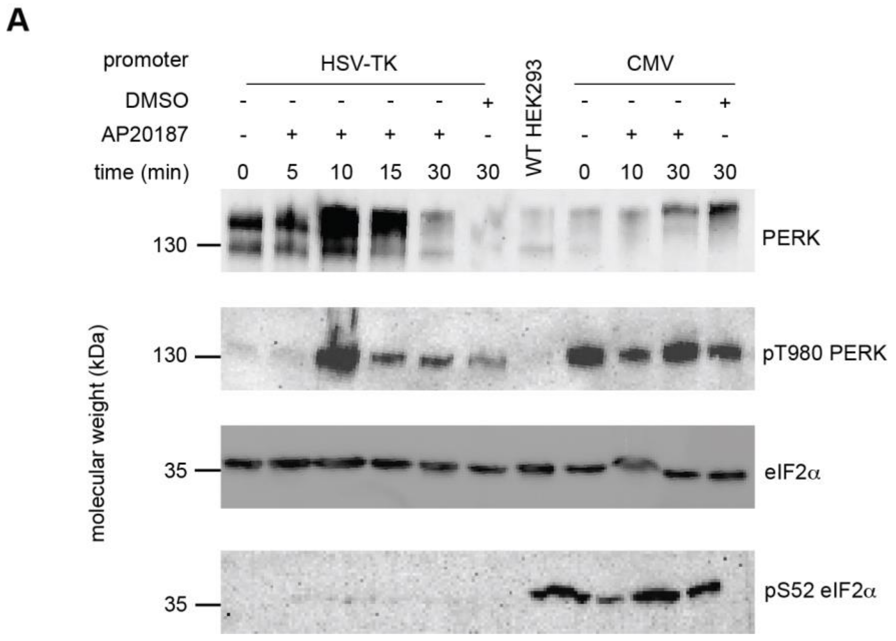

A. Biological replicate of experiment reported in Figure 3. Western blot analysis of dimerization-induced autophosphorylation. Blots against PERK, pT980 PERK, eIF2α and pS52 eIF2α are shown. Additionally, wild-type untransfected HEK293 cells were loaded alongside the two expression promoters. DMSO: solvent control, AP20187: dimerizer, HSV-TK: low-expression promoter, CMV: high-expression promoter.

#### Figure S8. Dimerization-dependent *cis*-autophosphorylation is conserved in the UPR and ISR kinases.

**A**

| Protein | Residue | Sequence window | Localization probability | Modification multiplicity | Spectral count | |
|---|---|---|---|---|---|---|
| **Gcn2** (post-auto-phosphorylation) | T882 | SSDNLTSAIGT | 82.1 % | 1 | 1 | canonical autophosphorylation site(s) |

**B**

| Protein | Residue | Sequence window | Localization probability | Modification multiplicity | Spectral count | |
|---|---|---|---|---|---|---|
| **HRI** (post-auto-phosphorylation) | tag | GSGGGSGGSAW | 90.4% | 1 | 7 | |
| | Y405 | KRGREYVDESA | 93.9 % | 1 | 6 | |
| | S408 | EYVDESACPYV | 91.5 % | 1 | 15 | |
| | T487 | RTPTHTSRVGT | 91.0 % | 1 | 8 | canonical autophosphorylation site(s) |
| | T542 | LTGLRTGQLPE | 92.8 % | 1 | 10 | |
| | FKBP | GKKVDSSRDRN | 91.2 % | 1 | 11 | |
| | FKBP | YAYGATGHPGI | 92.4 % | 1 | 27 | |

(spectral counts below 6 not shown)

**C**

| Protein | Residue | Sequence window | Localization probability | Modification multiplicity | Spectral count |
|---|---|---|---|---|---|
| **Ire1** (pre-auto-phosphorylation) | tag | GSGGGSGGSAW | 89.4% | 1 | 1 |
| | S806 | QNILVSTSSRF | 96.7 % | 1 | 3 |
| | S840 | LDSGQSSFRTN | 94.7 % | 1 | 3 |
| | S850 | NLNNPSGTSGW | 97.0 % | 1 | 35 |
| | S865 | ELLEESNNLQC | 97.3 % | 1 | 1 |
| | S876 | ETEHSSSRHTV | 92.1 % | 1 | 2 |
| | S1006 | NRDPPSALLMK | 96.9 % | 1 | 1 |

**D**

| Protein | Residue | Sequence window | Localization probability | Modification multiplicity | Spectral count: kinase-dead band | Spectral count: kinase-active band | |
|---|---|---|---|---|---|---|---|
| **Ire1** (post-auto-phosphorylation) | S806 | QNILVSTSSRF | 95.5 % | 1 | 2 | 3 | |
| | S840 | LDSGQSSFRTN | 96.7 % | 1;2 | 10 | 17 | canonical autophosphorylation site(s) |
| | S841 | DSGQSSFRTNL | 96.5% | 1;2;3 | 8 | 12 | canonical autophosphorylation site(s) |
| | T844 | QSSFRTNLNNP | 95.7% | 1;2;3 | 7 | 12 | canonical autophosphorylation site(s) |
| | S850 | NLNNPSGTSGW | 97.2 % | 1;2;3 | 2 | 11 | |
| | T881 | SSSRHTVVSSD | 96.2 % | 1 | 3 | 3 | |
| | T885 | HTVVSSDSFYD | 93.9% | 1 | 0 | 3 | |
| | S887 | VVSSDSFYDPF | 96.1 % | 1 | 3 | 3 | |

(spectral counts below 3 not shown)

A. Tandem mass spectrometry analysis of Gcn2 autophosphorylation after 60 minutes (corresponding to intact mass spectrum of the sample at 60 min reported in Figure 5C).

B. Tandem mass spectrometry analysis of HRI autophosphorylation after 120 minutes (corresponding to intact mass spectrum of the sample at 120 min reported in Figure 5F).
C. Tandem mass spectrometry analysis of Ire1 after purification. S850 is not a canonical autophosphorylation-site in the Ire1 activation loop[2] (corresponding to intact mass spectrum of the sample at 0 min reported in Figure 5I).
D. Tandem mass spectrometry analysis of Ire1 autophosphorylation after 60 minutes (corresponding to intact mass spectrum of the sample at 60 min reported in Figure 5I). Kinase-active and kinase-dead protomers were distinguished by SDS-PAGE gel electrophoresis and the respective bands were independently analyzed by tandem mass spectrometry.

#### Supplementary Table 1: Description of constructs used in this study.

| Construct name | *Residue range* and mutations | Description | Organism |
|---|---|---|---|
| PERK$^{KD}$-FKBP | *580-1082* Δ700-869 S675G T676G S681G S682G S684G S690G | M-6H-PERK-3C-4G-FKBP | *Mus musculus* |
| PERK$^{KD}$ inactive (heterodimer) | | M-6H-PERK(D935N)-6G-FKBP | |
| PERK$^{KD}$ active (heterodimer) | | M-TwinStrep- PERK-3C-4G-FKBP | |
| PERK$^{KD}$-FKBP (D588N) | | M-6H-PERK(D588N)-3C-4G-FKBP | |
| PERK$^{KD}$-FKBP (V610G) | | M-6H-PERK(V610G)-3C-4G-FKBP | |
| PERK$^{KD}$ (construct from Cui 2011) | *580-1082* Δ702-866 | M-6H-PERK | |
| CMV PERK$^{FL}$-3C-FKBP12$^{F36V}$-TEV-ALFA | *1-1114* | PERK-3C-4G-FKBP12F36V-TEV-4G-P-ALFA | |
| HSV-TK PERK$^{FL}$-3C-FKBP12$^{F36V}$-TEV-ALFA | | | |
| eIF2α | *1-315* | eIF2alpha-TEV-6G-GST | |
| Gcn2$^{KD}$ inactive (heterodimer) | *592-982* Δ666-767 | M-6H-Gcn2(D835N)-6G-FKBP | *Saccharomyces cerevisiae* |
| Gcn2$^{KD}$ active (heterodimer) | | M-TwinStrep- Gcn2-3C-4G-FKBP | |
| HRI$^{KD}$ inactive (heterodimer) | *160-585* Δ243-372 (replaced with 2G) | M-6H-HRI(D442N)-6G-FKBP | *Homo sapiens* |
| HRI$^{KD}$ active (heterodimer) | | M-TwinStrep- HRI-3C-4G-FKBP | |
| Ire1$^{KD\text{-}RNase}$ inactive (heterodimer) | *641-1115* | M-6H-Ire1(D797N)-6G-FKBP | *Saccharomyces cerevisiae* |

| | | |
|---|---|---|
| Ire1$^{KD\text{-}RNase}$ active (heterodimer) | | M-TwinStrep- Ire1-3C-4G-FKBP |

#### Supplementary Table 2: Intact mass spectrometry of heterodimers.

Expected and experimental masses for all constructs determined by intact mass spectrometry.

| Kinase | protomer | Unphosphorylated/phosphorylated | Expected MW [0P species] (Da) | Experimental molecular weight (Da) | | | | |
|---|---|---|---|---|---|---|---|---|
| | | | | 0P | 1P | 2P | 3P | 9P |
| PERK | Dead | Unphosphorylated | 52348.44 | 52348.70 | | | | |
| | active | | 51427.52 | 51427.93 | | | | |
| | Dead | phosphorylated | 50599.55 | 50599.60 | | | | |
| | active | | 39288.76 | 39290.72 | 39370.98 | | | |
| Gcn2 | Dead | Unphosphorylated | 45753.94 | 45754.03 | | | | |
| | active | | 46467.80 | 46468.18 | | | | |
| | Dead | phosphorylated | 45753.94 | 45753.97 | | | | |
| | active | | 46467.80 | 46468.18 | 46547.81 | | | |
| HRI | Dead | Unphosphorylated | 46486.76 | 46485.62 | | | | |
| | active | | 47199.64 | 47199.77 | | | | |
| | Active + tag | | | 51021.91 | | | | |
| | Dead | phosphorylated | 46486.76 | 46486.32 | | | | |
| | active | | 47199.64 | 47200.57 | 47279.64 | 47357.84 | | |
| | Active + tag | | 51152.82 | 51023.50 | 51101.09 | 51182.48 | 51262.22 | |
| Ire1 | Dead | Unphosphorylated | 66871.83 | 66872.59 | | | | |
| | active | | 67585.69 | 67586.24 | 67665.74 | | | |
| | Dead | phosphorylated | 66871.83 | | | 67032.60 | 67111.50 | |
| | active | | 67585.69 | | 67666.90 | …. | …… | 68304.70 |

### Supplementary Materials and Methods

#### Plasmids and cloning

All recombinant protein constructs are described in Supplementary Table 1.

Genes of all constructs were cloned from *Mus musculus* or *Homo sapiens* cDNA and *Saccharomyces cerevisiae* gDNA using KOD polymerase and touchdown PCR. All inserts were cloned into their respective backbones by Gibson assembly (Gibson MasterMix: T5 exonuclease 1U/µL, Phusion polymerase 2U/µL, Taq ligase 40U/µL in buffer containing 1mM dNTPs, 25% PEG-8000, 50 mM $MgCl_2$, 500 mM Tris pH 7.5, 5 mM NAD, 50 mM DTT, made in-house). Mutations were introduced using site-directed mutagenesis with Gibson assembly (NEBuilder HiFi Gibson assembly MasterMix). Constructs were transformed into chemically competent *E. coli* OMNIMAX, and plasmid DNA was purified using QIAprep 2.0 Spin Miniprep Columns (Qiagen). Plasmids were sequenced by Microsynth.

#### Recombinant protein expression

Sequences for recombinant protein expression were derived from *Mus musculus* unless otherwise specified. All sequences and construct boundaries are summarized in Supplementary Table 1. All recombinant proteins were expressed in *E. coli* Rosetta2 cells. All kinase domain-containing proteins were co-expressed with lambda phosphatase. Pre-cultures of *E. coli* Rosetta2 were grown in lysogeny broth (LB) overnight before inoculating the main culture to an OD of 0.05. Large scale cultures of *E. coli* Rosetta2 were grown in TB media supplemented with 1% w/v glucose to an OD of 1.5 before addition of 200 nM IPTG and expression overnight at 20°C.

#### Recombinant protein purification

##### *PERK$^{KD}$*

Cell pellets from 1L culture were lysed at 4°C with stirring in 50 mL buffer NiNTA-A (150 mM NaCl, 50 mM Tris pH 7.4, 1 mM TCEP) supplemented with 1 mM $MnCl_2$, 1 mM $MgCl_2$, 2 mM benzamidine, 1 µg/mL benzonase (in-house) and 10 mg lysozyme. The lysate was sonicated on a Branson W-450D Sonifier for 10 minutes at 50% amplitude, and clarified at 38,000 g, 4°C, 1 h.

Cleared lysate was supplemented with 20 mM imidazole and loaded onto a 5 mL HisTrap HP (Cytiva) column using the AEKTA Pure liquid chromatography system. The column was washed with 100 mL NiNTA-A and 20 mL NiNTA-A with 20 mM imidazole. Stepwise elution was performed with increasing concentrations of imidazole. 100 mM and 300 mM elution fractions were pooled, diluted to 100 mM imidazole and dialyzed overnight in a SpectraPor 8-12 kDa cutoff dialysis tube (Repligen) with 1 mM $MnCl_2$ and 1 mL of 1 mg/mL TEV protease (in house) per 25 mL eluate.

Dialyzed eluates were diluted to 50 mM NaCl with Q-A buffer (50 mM Tris pH 7.4, 1 mM TCEP) and loaded onto a 1 mL HiTrap Q HP column (Cytiva) connected in series with a 5 mL HisTrap HP to remove uncleaved protein and His-tagged lambda phosphatase. The Q column was washed with 20 column volumes Q-A and $PERK^{KD}$ eluted with a 0-100% NaCl gradient in Q-B (1M NaCl, 50 mM Tris pH 7.4, 1 mM TCEP). Fractions containing $PERK^{KD}$ were pooled and concentrated using a 10 kDa cutoff Amicon filter for injection onto a Superdex 200 10/300 column equilibrated in NiNTA-A buffer. $PERK^{KD}$-containing fractions were pooled and concentrated with a 10 kDa cutoff Amicon filter before plunge-freezing and storage of 10 uL aliquots at -70°C.

##### *$PERK^{KD}$-FKBP*

Dimers of $PERK^{KD}$-FKBP were purified according to the protocol for $PERK^{KD}$, except for the addition of 1 mg AP20187 (Sigma Aldrich SML2838) dissolved in 200 μL of 96% ethanol to the clarified lysate. The lysate was incubated for 30 minutes in the cold room with AP20187 before loading it onto the HisTrap column.

##### *eIF2α*

Cell pellets from 1 L culture were lysed, sonicated and clarified under the same conditions as $PERK^{KD}$. Clarified lysate was incubated with 5 mL glutathione Sepharose 4B beads pre-equilibrated in NiNTA-A buffer. Beads were incubated for 3 h with rotation at 4°C, then centrifuged for 30 min, 4°C, 500 g. Beads were washed three times with NiNTA-A. Beads were incubated with 1 mL of 1 mg/mL TEV protease (in-house) per 25 mL cleavage volume overnight. The supernatant was diluted to 50 mM NaCl with Q-A buffer and loaded onto a 1 mL HiTrap Q HP column (Cytiva). The column was washed with 20 CV Q-A and eIF2α was eluted with a 0-

100% NaCl gradient in Q-B. Fractions containing eIF2$\alpha$ were pooled and concentrated using a 10 kDa cutoff Amicon filter for injection onto a Superdex 200 10/300 column equilibrated in NiNTA-A buffer. eIF2$\alpha$-containing fractions were pooled and concentrated with a 10 kDa cutoff Amicon filter before plunge-freezing and storage of 10 uL aliquots at -70°C.

##### *PERK, Gcn2, HRI, and Ire1 heterodimers*

Purification of all heterodimers followed the same, sequential protocol: (1) HisTrap (2) StrepTrap (3) ion exchange (4) size-exclusion. For each heterodimer, two expression cultures were grown: one of the active, TwinStrep-tagged kinase and one of the kinase-inactive, $His_6$-tagged kinase construct, both co-expressed with lambda phosphatase. The two pellets were lysed together, sonicated and the lysate clarified. AP20187 was then added to the cleared lysate. The lysate was loaded on a HisTrap column, washed with NiNTA-A and 2% NiNTA-B. Proteins were eluted with 200 mM imidazole directly onto a StrepTactin XT column (Cytiva) equilibrated in NiNTA-A. Following elution, the HisTrap column was removed, and the StrepTactin column washed with NiNTA-A. The protein was eluted from the StrepTactin column with 20 mM biotin in NiNTA-A, pH 7.4. Heterodimer-containing fractions were pooled and 1 mL of 1 mg/mL TEV (in house) was added per 25 mL eluate. For PERK heterodimer purification, the cleaved protein was flowed over HisTrap and Q columns connected in series and heterodimer-containing fractions were concentrated and injected onto a S200 10 300 column equilibrated in NiNTA-A buffer. Gcn 2 heterodimers were purified as described above, except that a HiTrap Capto S column (Cytiva) was used for ion exchange. Ire1 heterodimers were purified as described above, except that a HiTrap Heparin HP (Cytiva) column was used in place of ion exchange.

#### Mass spectrometry

##### *Intact protein liquid chromatography-mass spectrometry*

LC-MS analysis was performed on a Vanquish™ Horizon UHPLC system (Thermo Scientific) coupled to a Synapt G2-Si mass spectrometer (Waters) equipped with a ZSpray ESI source (Waters). Up to 1 pmol of protein in 2 µL solution were loaded to a XBridge Protein BEH C4 column (300Å, 2.5 µm particle size, dimensions 2.1 mm x 150 mm; Waters). On a BioResolve Polyphenyl mAb (450Å, 2.7 µm, 2.1 mm X 150 mm; Waters) with a working temperature of 50°C, 0.1% formic acid (FA) as solvent A, 100% acetonitrile (ACN), 0.08% FA as solvent B. Proteins were separated on a 6 min step gradient from 12% to 72% solvent B respectively from 15% to 80% at a flow rate of 250 µL/min. The mass spectrometer was operated in resolution mode, the capillary voltage set to 2 kV, a source temperature of 120°C, and a desolvation temperature of 400°C. Data were recorded with MassLynx V 4.2 (Waters) and calibrated with the lockmass of Glu-Fibrinopeptide B. Raw data were analyzed using the MaxEnt 1 process to reconstruct the uncharged average protein mass.

##### *Bottom-up liquid chromatography-mass spectrometry*

###### *Sample preparation*

Coomassie-stained gel bands were destained with a mixture of acetonitrile and 50 mM ammonium bicarbonate (ABC). The proteins were reduced with 10 mM dithiothreitol and alkylated with 50 mM iodoacetamide. Trypsin (Promega; Trypsin Gold, Mass Spectrometry Grade) was used for proteolytic cleavage. Digestion was carried out at 37°C overnight. Formic acid (10%) was used to stop digestion and 5% FA, 50% ACN was used to extract peptides from the gel.

Proteins in buffers containing CHAPS were precipitated by adding a 4-fold volume of ice-cold acetone and kept overnight at -20°C or for 3h at 4°C. Proteins were pelleted by centrifugation for 10 min at 15000 g. Supernatant was discarded and pellets were washed with ice-cold 80% acetone and centrifugated 10 min at 15000 g. The wash step was omitted if the precipitated protein amount was lower than 1 µg. After removing the supernatant, tubes were left open to dry the pellets.

Protein pellets (1-4 µg) were dissolved in 10 µL 8 M urea, 50 mM ABC, vortexed and briefly sonicated. Dithiothreitol (DTT, 250 mM stock solution in dH2O) was added to a final concentration of 10 mM and let react for 30 minutes at RT. Reduced thiols were alkylated by adding iodoacetamide (500 mM stock solution in dH2O) and let react for 30 min at RT in the dark. Residual iodoacetamide was quenched with half the amount of DTT used in the reduction step. Prior to adding proteases, the solution was diluted to 1 M urea with 50 mM ABC. Proteases were added at a 1:30 (protease:protein) ratio and incubated at 37°C overnight (for trypsin) or at 25°C for 2-5 h for chymotrypsin. Digestion was stopped by acidifying the solution with 10% TFA. Peptides were purified on StageTips[3].

Digest clean-up protocol is based on Rappsilber et al., 2007[3] with some modifications. Three C18 Empore disk punches were packed into 200 µL pipette tips. All applied solutions were passed through the tip by centrifugation. Tips were prewetted with 100 µL methanol, followed by 100 µL 80% acetonitrile (ACN), 0,1% TFA, and equilibrated with 100 µL 0.1% TFA by spinning at 376 g for 4 - 6 min. Acidified peptide solutions were applied, and spun at 271 g. Bound peptides were washed with 100 µL 0.1% TFA, and eluted with 2 x 30 µL 40% ACN, 0.1% TFA into PCR tubes. Eluates were dried in a vacuum centrifuge and resuspended in 20 µL 2% ACN, 0.1% TFA.

##### *Liquid chromatography-mass spectrometry analysis*

LC-MS analysis was performed on a Vanquish Neo UHPLC system (Thermo Scientific) coupled to an Orbitrap Exploris 480 mass spectrometer (Thermo Scientific). The system was equipped with a Nanospray Flex ion source (Thermo Scientific), and a Column Heater (IonOpticks) connected to a Heater Controller (IonOpticks).

Peptides were loaded onto a trap column (PepMap Neo C18 5mm × 300 µm, 5 µm particle size, Thermo Scientific) using 0.1% TFA as mobile phase, and separated on an analytical column (e.g. Aurora Ultimate XT C18, 25 cm × 75 µm, 1.7 µm particle size, IonOpticks), applying a linear gradient starting with a mobile phase of 98% solvent A (0.1% formic acid (FA)) and 2% solvent B (80% acetonitrile, 0.08% FA), increasing to 35% solvent B over 30 min at a flow rate of 300 nL/min. The analytical column was heated to 50°C. The mass spectrometer was operated in data-dependent acquisition (DDA) mode, with 1.2 s MS1 cycle time. Survey scans were acquired from 375-1500 m/z with lock mass enabled, normalized AGC target of 100%, resolution of 60,000. The most intense precursor ions (charge states +2 to +6) were selected for fragmentation using an isolation window of 1.4 m/z. Selected ions were analyzed with a maximum fill time of 100 ms, normalized AGC target of 100%, and resolution of 30,000 after HCD fragmentation with normalized collision energy of 30%. Monoisotopic precursor selection (MIPS) was set to “peptide” mode, the intensity threshold to $2.5 \times 10^4$, and selected precursors were dynamically excluded for 10 seconds with isotope exclusion enabled.

##### *Mass spectrometry data analysis*

MS raw data were analyzed with FragPipe, using MSFragger[4], IonQuant[5], and Philosopher[6]. The default FragPipe workflow for label free quantification was used, except "Normalize intensity across runs" and "Match between runs" were turned off. Cleavage specificity was set to the respective protease with 2-4 missed cleavages allowed. The false discovery rate (FDR) was set to 1%. Oxidation of methionine, phosphorylation on serine, threonine and tyrosine and N-terminal protein acetylation were specified as variable modifications. Carbamidomethylation of cysteine was set as a fixed modification. MS2 spectra were searched against target sequences in their expression background (*E.coli*), concatenated with a database of common laboratory contaminants (https://github.com/maxperutzlabs-ms/perutz-ms-contaminants).

Computational analysis was performed using Python and the Python library MsReport (source code: https://github.com/hollenstein/msreport)[7].

#### Mass photometry

Protein samples were centrifuged at 21,000 g, 4°C, 5 minutes after thawing. Samples were diluted to 500 nM in 150 mM NaCl, 50 mM Tris pH 7.4, 1 mM TCEP. Microscopy coverslips were cleaned sequentially by sonication in water, isopropanol, and then water again and dried under constant air flow. All measurements were recorded on a TwoMP mass photometer (Refeyn) for 60 seconds at the indicated final protein concentration. Protein mass was calculated using a contrast-to-mass calibration with either NativeMark Protein Standard (Invitrogen) or MassFerence P2 standard (Refeyn). Data analysis was performed using Discover MP (Refeyn).

#### Circular dichroism

Monomeric $PERK^{KD}$-FKBP, $PERK^{KD}$-FKBP (D588N), and $PERK^{KD}$-FKBP (V610G) were buffered into 20 mM $NaH_2PO_4$/NaOH pH 7.4 using Zeba™ Spin Desalting columns 7K MWCO (Thermo Scientific). Spectra were recorded using a Chirascan Plus Spectrophotometer (Applied Photonics) at 20°C in a 0.1 cm quartz cuvette from 180 to 340 nm wavelength, with simultaneous absorbance measurements at 280 nm wavelength. Three scans were averaged for each sample, and the corresponding buffer spectrum was subtracted from the final spectrum. Final cuvette concentration was calculated based on absorbance at 280 nm.

#### Nano Differential Scanning Fluorimetry (nDSF)

nDSF measurements were performed on a Prometheus NT.48 (NanoTemper Technologies). 0.5 mg/mL of monomeric $PERK^{KD}$-FKBP, $PERK^{KD}$-FKBP (D588N), and $PERK^{KD}$-FKBP (V610G) were loaded into NanoTemper NT.48 standard capillaries in triplicates. Intrinsic fluorescence (350/330 nm) was monitored during a linear temperature gradient from 20 to 95°C at a heating rate of 1°C/min. Melting temperatures ($T_m$) were determined from the first derivative of the fluorescence ratio.

#### Bissulfosuccinimidyl suberate (BS3) protein crosslinking

Monomeric $PERK^{KD}$-FKBP, $PERK^{KD}$-FKBP (D588N), and $PERK^{KD}$-FKBP (V610G) were buffer exchanged into 50 mM HEPES pH 7.4, 150 mM NaCl, 1 mM DTT, 1mM ATPγS, 2 mM $MgCl_2$ using Zeba™ Spin Desalting columns 7K MWCO (thermo scientific). Proteins were crosslinked at 15 µM in the absence of BS3 (1:0), or with 1.5 µM (1:10) and 0.15 µM (1:100) BS3 for 30 minutes and quenched in Tris pH 7.4. Samples were analyzed using 12% SDS-PAGE gels.
